# Sleep fragmentation drives local, network-specific epileptic activity in the human brain

**DOI:** 10.64898/2025.11.30.691386

**Authors:** Birgit Frauscher, Alyssa Ho, Barbora Matoušková, Kassem Jaber, Tamir Avigdor, Erica Minato, John Thomas, Derek Southwell, Jeffery Hall, David Carlson, Petr Klimes, Laure Peter-Derex, Sana Hannan

## Abstract

Sleep has complex links with epileptic activity, yet the causal role of sleep instability in driving and modulating pathological discharges in the human brain remains incompletely understood. Here we directly examine this by characterising the fine-scale temporal coupling between experimentally induced sleep arousals and interictal epileptiform discharges (IEDs), using combined stereo-electroencephalography and polysomnography recordings in patients with epilepsy. Sleep arousals triggered rapid IED increases, with effects gated by anatomical region and sleep stage. Increases were confined to neocortical regions and occurred during both non-rapid eye movement stage 2 (N2) and stage 3 (N3) sleep, with a larger effect observed in N2. IED increases did not differ between the seizure-onset zone and surrounding regions. Despite elevating IED counts, arousals did not alter IED spatial propagation, indicating state-dependent enhancement of local cortical excitability without recruitment of broader epileptic networks. These findings establish a causal role for sleep instability in actively driving pathological activity on fine-grained spatiotemporal scales, and highlight sleep stabilisation as a promising therapeutic strategy to reduce epileptic burden and preserve cortical network function.

## Introduction

More than half of individuals with epilepsy experience sleep problems, including poor sleep quality, insomnia and excessive daytime sleepiness.^1^ Sleep disturbances not only impair quality of life but can also promote seizures, creating a vicious cycle that further exacerbates the epileptic state.^2,3^ Nocturnal seizures further increase the risk of sudden unexpected death in epilepsy (SUDEP), the leading cause of epilepsy-related mortality.^4^ Previous work has established a close relationship between sleep instability and epileptic activity. Seizure susceptibility during sleep-wake transitions is well recognised, from early electroclinical observations including both primary generalised and focal epilepsy syndromes to more recent studies showing that epileptic activity preferentially occurs during non-rapid eye movement (NREM) sleep instability.^5–11^ Sleep arousals are characterised by transient increases in vigilance and abrupt EEG frequency shifts lasting 3-15 s, and represent intermediate states between sleep and wakefulness which plays a critical role in sleep-wake regulation.^8,12,13^ The cycling alternating pattern (CAP), a marker of sleep instability which consists of EEG arousal-related transients, has been linked to epileptic activity.^8,14–16^ In addition, arousal-related events such as those induced by sleep-disordered breathing have been associated with increased seizure occurrence.^14,17^ Intracranial studies using stereo-electroencephalography (SEEG) have further demonstrated that epileptic activity is temporally linked to arousal fluctuations,^18,19^ with rates of interictal epileptiform discharges (IEDs) increasing prior to arousal onset and, in neocortical regions, during the arousal itself.^19^ More recent mechanistic work supports the idea that unstable slow oscillations and arousal-related dynamics facilitate epileptic activity,^20^ and that this activity occurs more frequently during lighter, less stable NREM stage 2 sleep (N2) than during deeper, more stable NREM stage 3 sleep (N3).^9,20,21^

Despite these studies, several important gaps remain. First, much of the existing literature has relied on observational designs, making it difficult to disentangle causal relationships between sleep instability and epileptic activity – specifically, whether one influences the other directly or both are driven by a separate shared mechanism. Secondly, many studies have relied on scalp electroencephalography (EEG), which often fails to detect focal or deep-seated epileptic activity.^3,22–24^ Consequently, there is a need for experimental studies to explore causality through direct manipulation of sleep-related neural networks in the human epileptic brain.

Stereo-electroencephalography (SEEG), as performed in patients with drug-resistant epilepsy undergoing presurgical evaluation, provides unique access to deep brain structures with high spatiotemporal resolution.^23,24,29^ When combined with polysomnography (PSG) – including scalp EEG, electromyography (EMG), and electrooculography (EOG) – SEEG enables detailed characterisation of the temporal dynamics between sleep stability and epileptic activity.^11,23,30^

Here, using combined SEEG-PSG recordings from patients with drug-resistant focal epilepsy undergoing presurgical evaluation, we experimentally manipulated sleep stability to assess its direct impact on epileptic activity. We used auditory stimulation to induce sleep arousals, thereby creating controlled, transient perturbations of sleep instability,^31^ and compared the counts and propagation of IEDs before and during these events. This perturbational approach enabled a more direct test of the immediate effects of arousal-related sleep instability on IED occurrence in the human brain. Focusing on the interictal network allowed high temporal specificity and sensitivity to detect immediate changes in network excitability, as IEDs can be precisely time-locked to external events, offering a tractable window into state-dependent pathological activity. Importantly, IEDs are associated with cognitive disruption and impaired memory consolidation.^32–36^ Given the important role of sleep in these processes,^31,37,38^ understanding how arousals influence IED expression provides mechanistic insight into how sleep fragmentation may worsen cognitive outcomes in epilepsy, even in the absence of seizures.

We hypothesised that arousals would increase epileptic spiking and promote more widespread propagation, reflecting heightened network excitability. Consistent with this hypothesis, we observed a significant increase in IED counts immediately following arousal onset compared to pre-arousal periods, indicating a direct influence of arousals on epileptic dynamics. We further examined whether this effect was modulated by brain region, seizure-onset zone (SOZ) involvement, and sleep stage. We predicted greater arousal-related spiking in neocortical than mesiotemporal regions, based on stronger functional connectivity between the neocortex and thalamus, which plays a central role in sleep regulation,^39^ and a larger effect during N2 sleep, where coupling between arousals and epileptic activity is strongest.^19^

Consistent with our hypotheses, our findings revealed an increase in IED counts in the neocortex and during both N2 and N3 sleep, with a larger effect in N2, highlighting the critical role of thalamocortical circuits and sleep-stage dynamics in modulating interactions between arousals and epileptic activity. Arousal-related IED increases did not differ between the SOZ and surrounding regions, suggesting that the SOZ is not preferentially engaged by transient arousal-induced excitatory changes. In contrast to the effects on IED counts, arousals did not influence the spatial propagation of IEDs. Together, our findings suggest that transient sleep arousals selectively enhance local cortical excitability without engaging broader network dynamics required for IED propagation, and that this relationship is dependent on brain region and sleep stage. This work therefore provides direct evidence linking evoked sleep arousals to local modulation of epileptic activity, helping to refine current models of how transient fluctuations in brain state modulate epileptic dynamics and highlighting sleep stability as a key determinant of pathological activity in the human brain.

## Results

### Auditory stimulation induces sleep fragmentation and arousals in patients with epilepsy

To examine the impact of evoked sleep arousals on interictal activity, we first verified that our auditory stimulation protocol reliably induced arousals and sleep fragmentation in patients with drug-resistant focal epilepsy undergoing overnight SEEG-PSG recordings (*n* = 17). Auditory tones of varying intensities were delivered during stable NREM sleep in the first four hours of the night, using a validated approach designed to increase arousal counts without inducing full awakenings, guided by real-time PSG-based sleep scoring (**Fig. 1**).^13,31^ Across participants, total sleep time (TST) was 400.9 ± 89.9 min (mean ± SD, range 214.5-542.0 min) (**Fig. 2a**, see **Supplementary Table 1** for clinical and demographic details). Auditory stimulation induced 440 evoked arousals in total, corresponding to 25.9 ± 9.8 arousals per participant (range 9-48) (**Fig. 2b**). A mean of 31.6 ± 17.2% of auditory stimuli (range 10.7-86.2%) successfully triggered arousals (**Fig. 2c**), with 90 dB being the most used stimulus volume (**Fig. 2d**), although responsiveness to stimulus intensity varied across participants (**Supplementary Fig. 1**). Evoked arousals had a mean duration of 7.8 ± 3.3 s (range 3-15 s) (**Supplementary Fig. 2a**), and occurred more frequently during N2 (14.7 ± 7.0, range 1-32) than N3 sleep epochs (5.2 ± 6.4, range 0-29) (**Fig. 2e**). Importantly, the overall arousal rate – including both evoked and spontaneous arousals occurring independently of auditory stimulation – was significantly higher during the first half of the night, when stimulation was delivered (27.0 ± 9.5 events/hour (median ± IQR), range 11.5-74.3 events/hour), compared to the second half (18.0 ± 10.2 events/hour, range 5.8-52.4 event/hour) (Wilcoxon signed rank test, *p* = 0.00118, Cliff’s *d* = 0.46, *n* = 17, **Fig. 2f**). Together, these results confirm that auditory stimulation effectively increased sleep fragmentation in this cohort.

**Figure 1.**
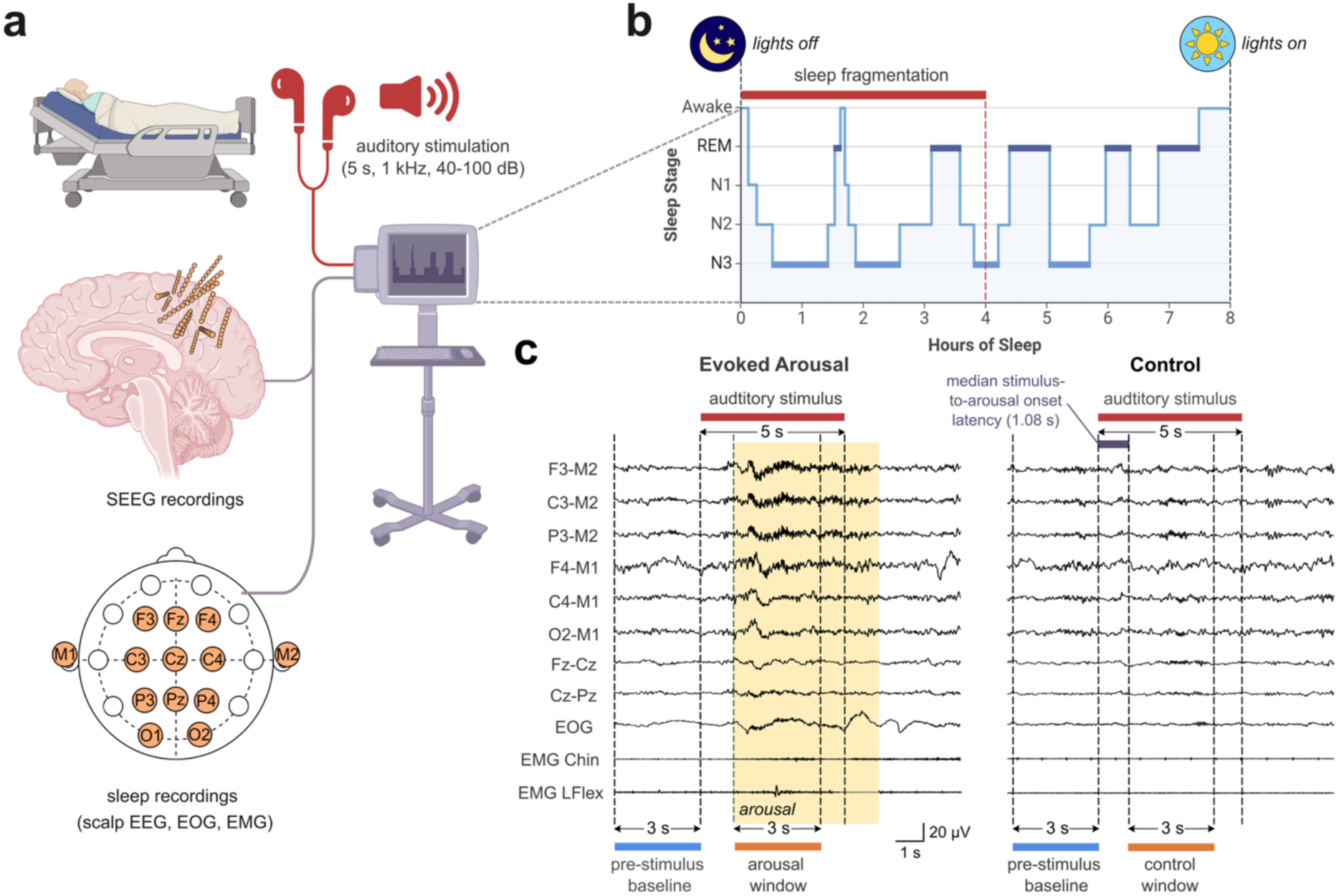
Overview of the sleep fragmentation protocol during overnight SEEG-PSG recordings. **(a)** Simultaneous SEEG and sleep (PSG) recordings were acquired during overnight sleep in patients with drug-resistant focal epilepsy undergoing their presurgical evaluation. SEEG electrodes were implanted according to each patient’s clinical requirements as part of their diagnostic workup. PSG included subdermal scalp EEG recordings positioned according to the standard international 10-20 system, alongside surface electrooculography (EOG) and electromyography (EMG) from the chin and tibialis anterior muscles (LFlex/RFlex). **(b)** Sleep fragmentation was induced in the first half of the sleep night, starting from the patient’s habitual bedtime, by delivering 5-second, 1 kHz auditory tones at intensities ranging from 40 to 100 dB during stable NREM sleep. Auditory stimulation was manually delivered through earphone inserts based on real-time PSG-based sleep scoring to induce arousals without full awakenings. **(c)** Arousals were annotated offline using PSG recordings; an event was classified as an ‘evoked arousal’ if it began within 6.5 s of stimulus onset. Interictal epileptiform discharge (IED) counts and propagation were compared across 3-second epochs: a pre-stimulus baseline and a post-stimulus onset window. For stimuli that evoked an arousal, the post-stimulus onset window began at arousal onset. For non-arousing stimuli, it began 1.08 s after stimulus onset, corresponding to the median stimulus-to-arousal onset latency (latency-matched control window).

**Figure 2.**
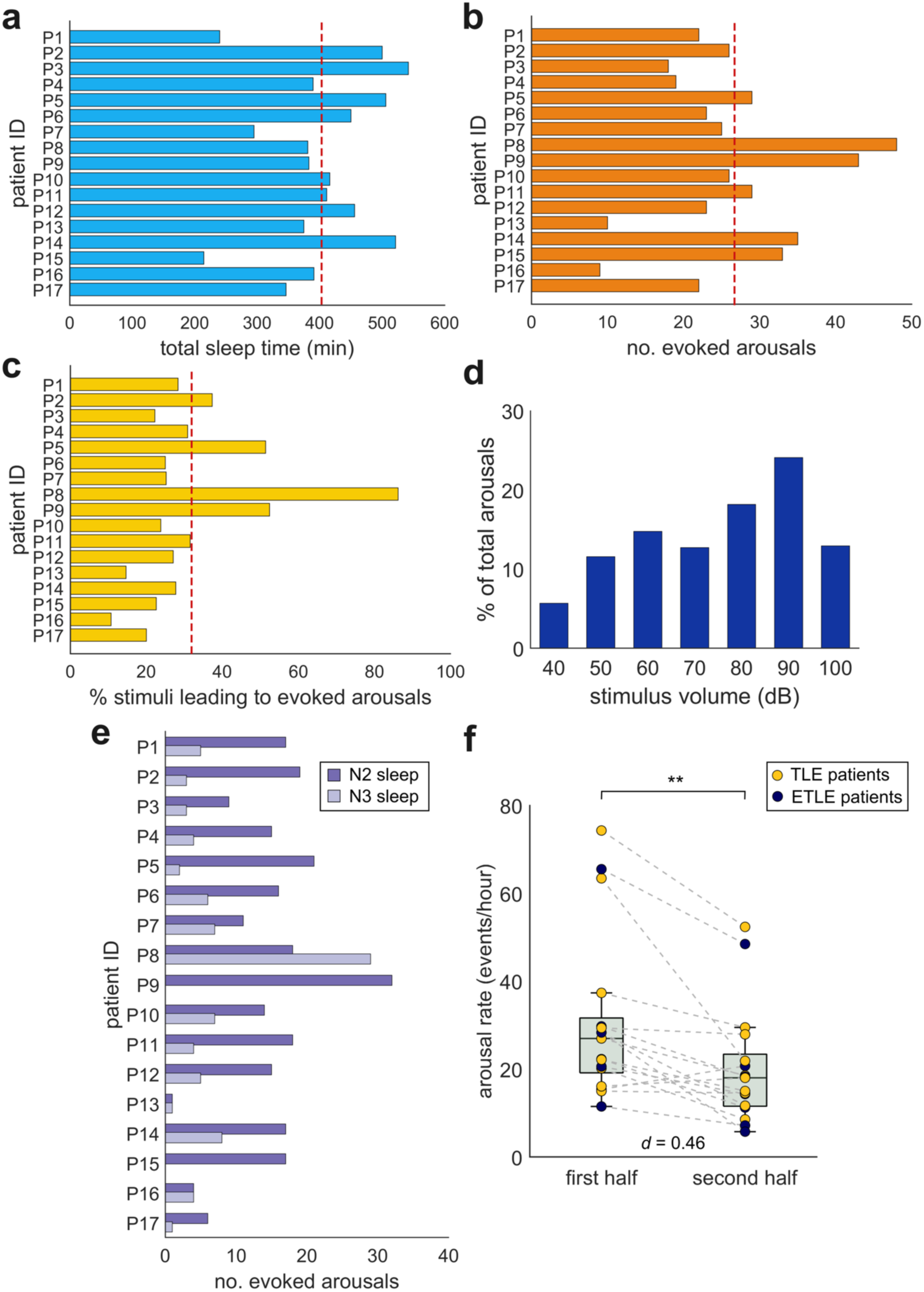
Auditory stimulation reliably induces evoked arousals and sleep fragmentation in patients with epilepsy. **(a)** Total sleep time (TST) during overnight SEEG-PSG recordings across all patients. **(b)** Total number of evoked arousals induced by the auditory stimulation protocol in each participant. **(c)** Percentage of stimuli that lead to evoked arousals in each participant. The red dashed line in panels (a-c) indicates the group mean. **(d)** Distribution of auditory stimulus volumes used to evoke arousals, pooled across all patients. 90 dB was the most commonly used volume for inducing evoked arousals. **(e)** Number of evoked arousals in each participant during epochs scored as NREM stage 2 (N2) and stage 3 (N3) sleep. Arousals overlapping stage transitions were excluded from stage-specific counts to ensure stable sleep state classification. **(f)** Total arousal rate, including both auditory evoked arousals and spontaneous arousals independent of auditory stimulation, during the first and second halves of the sleep night. A significantly greater number of arousals occurred during the first half of the night, when the auditory stimulation protocol was administered (Wilcoxon signed-rank test, *p* = 0.00118, Cliff’s *d* = 0.46). Data are presented as individual patient values. Patients with temporal lobe epilepsy (TLE; yellow circles) and extratemporal lobe epilepsy (ETLE; navy blue circles) are distinguished for visual comparison. No consistent differences in arousal rates between TLE and ETLE subgroups were observed across the first and second half of the night. These outcomes confirmed the effectiveness of the auditory stimulation protocol in patients with epilepsy.

### Evoked arousals acutely increase interictal epileptiform discharge counts but not their spatial propagation

We then asked whether transient sleep arousals triggered by auditory stimulation modulate epileptic activity. IEDs were detected on bipolar SEEG channels using a validated detector,^40^ previously applied in epilepsy and sleep studies.^11,30^ The median number of SEEG channels involved across IEDs used in the analysis was 2 (interquartile range, IQR 5; range 1-135) (**Supplementary Fig. 2b**). Global IED counts and rates per patient across the night are provided (**Supplementary Table 2).** IED count was defined as the number of unique IED events per epoch and, for each participant, IED counts were computed in 3-second epochs across four conditions: pre-stimulus and post-arousal onset windows for arousal-inducing stimuli, and matched pre-stimulus and latency-matched control windows for auditory stimuli that did not induce arousals (controls) (**Fig. 1c**). Contrasting evoked arousal and control conditions allowed us to isolate the specific effects of arousals on local interictal spiking, independent of auditory stimulation. To confirm that IED counts captured during pre-stimulus epochs were representative of each patient’s baseline, we compared IED counts between pre-stimulus epochs and randomly sampled arousal-free epochs across the night using a generalised linear mixed-effects mode (GLMM). IED counts did not differ significantly between the two epoch types (GLMM, β-estimate ± standard error (SE): -0.052 ± 0.027, *P* = 0.053, **Supplementary Fig. 3**), confirming that the 3-s pre-stimulus window was representative of baseline IED activity for each patient.

To determine whether interictal spiking varied across conditions, we modelled IED counts per channel-epoch using generalised linear mixed-effects modelling (GLMM), with condition (evoked arousal vs control) as the fixed effect of primary interest, adjusting for pre-stimulus baseline IED count and stimulus intensity, and including random intercepts for SEEG contact nested within patient (*n* = 17 patients, **Fig. 3**). This revealed a significant effect of condition, with predicted IED counts significantly higher during evoked arousals than controls (GLMM, β-estimate ± standard error (SE): 0.537 ± 0.012, incidence rate ratio (IRR) = 1.71, *p* < 0.0001), corresponding to a 71% increase in predicted IED count relative to controls (**Fig. 3d**). Exploratory subgroup stratifications by epilepsy type, MRI status, and MRI lesion category showed a consistent visual trend towards higher IED counts during arousals across all subgroups (**Supplementary Fig. 4**), suggesting the effect was not restricted to specific lesion or epilepsy subtypes. To rule out the possibility that the observed IED increase reflected a baseline brain state that was more permissive for arousal and independently associated with a higher IED rate, rather than an effect of the arousal itself, we compared IED counts in the 3-s pre-stimulus epochs preceding controls and evoked arousals and found no significant difference between conditions when adjusting for stimulus intensity (GLMM, β-estimate ± SE: 0.019 ± 0.014, IRR = 1.02, *P* = 0.182, **Supplementary Fig. 5**).

**Figure 3.**
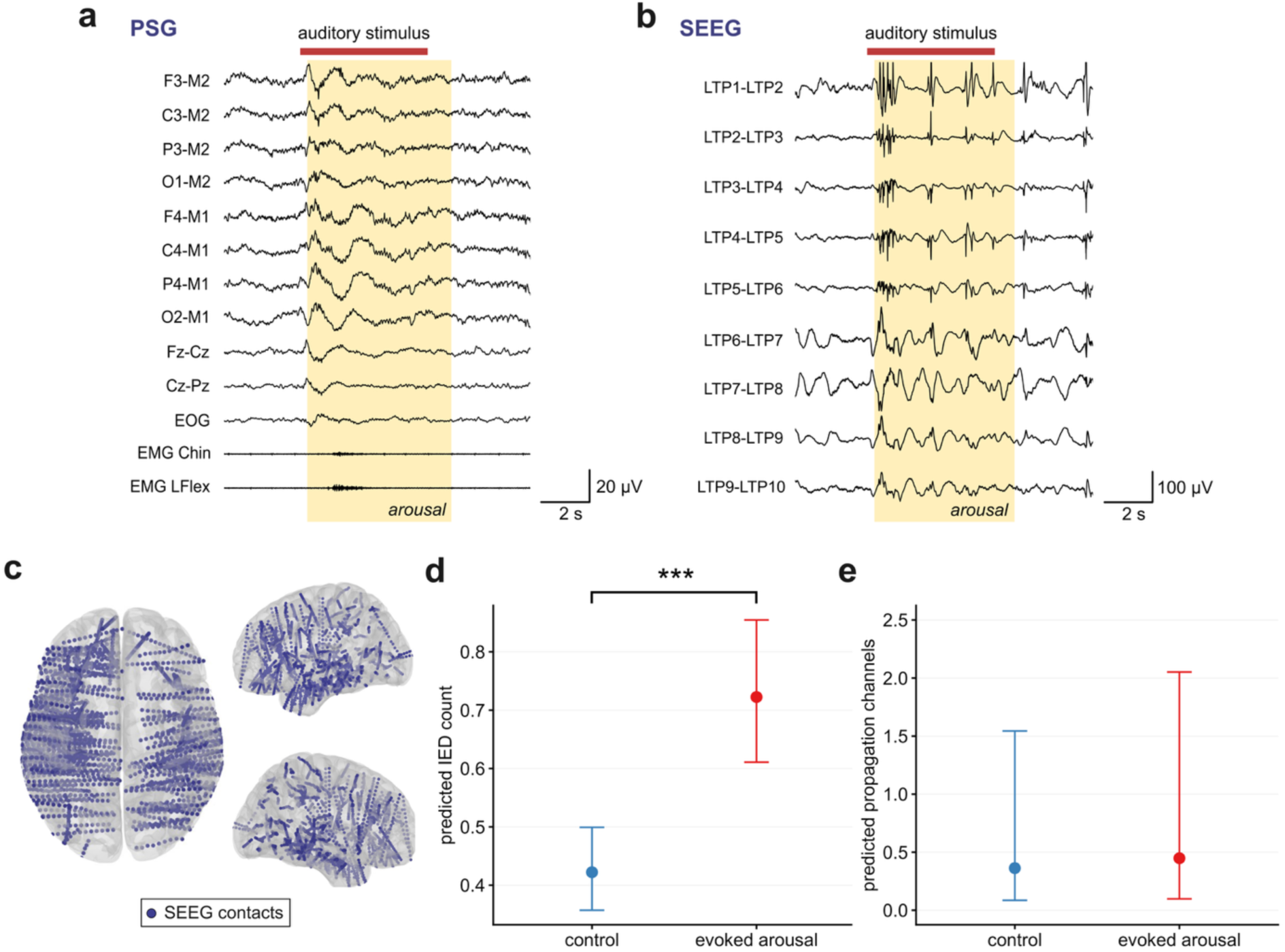
Evoked sleep arousals increase interictal epileptiform discharge (IED) counts but not propagation. **(a)** Representative example of polysomnography (PSG) traces from one patient showing scalp EEG, electrooculography (EOG) and electromyography (EMG) from chin and tibialis anterior (LFlex) channels. The yellow shaded region indicates an evoked arousal, induced by the auditory stimulus, indicated by the red bar. **(b)** Corresponding SEEG traces from the same time window from a depth electrode targeting the left temporal lobe (bipolar channels LTP1-LTP10). A marked increase in IEDs is observed during the arousal period (yellow shaded region), following auditory stimulation. **(c)** Spatial distribution of SEEG contacts across all patients (*n* = 17 patients), projected onto a standard brain template. **(d)** Model-predicted mean IED counts per channel-epoch under control and evoked arousal conditions, adjusted for pre-stimulus IED count and stimulus intensity. Predicted IED count was significantly higher during evoked arousals than controls (generalised linear mixed-effects model (GLMM), β-estimate ± standard error (SE): 0.537 ± 0.012, incidence rate ratio (IRR) = 1.71, *p* < 0.0001), corresponding to a 71% increase relative to controls. **(e)** Model-predicted IED propagation channel numbers per epoch under control and evoked arousal conditions, adjusted for pre-stimulus propagation channel number and stimulus intensity. The number of propagation channels did not differ significantly between evoked arousals and controls (GLMM, β-estimate ± SE: 0.212 ± 0.408, IRR = 1.24, *P* = 0.604). Error bars represent 95% confidence intervals. \*\*\**p* < 0.001.

Spontaneous arousals were also associated with a significant, though comparatively modest, increase in predicted IED count during the arousal period (GLMM, β-estimate ± SE: 0.046 ± 0.010 SE, IRR = 1.05, *p* < 0.0001, **Supplementary Fig. 6a**). We also examined whether evoked arousals similarly modulate high-frequency oscillations (HFOs) and found that predicted HFO count was indeed significantly higher during evoked arousals than controls (GLMM, β-estimate ± SE: 0.530 ± 0.014, IRR = 1.70, *p* < 0.0001, **Supplementary Fig. 7**), corresponding to a 70% increase and mirroring the pattern observed for IEDs. These results demonstrate that arousals acutely increase IEDs as well as HFOs, an effect that cannot be explained by auditory stimulation alone, showing a direct link between sleep microstructure and transient increases in epileptic activity.

We then examined whether evoked arousals influence not only IED counts but also their spatial propagation. Whereas the previous analysis focused on changes in local excitability by analysing IED counts, examining propagation allowed us to determine whether transient sleep arousals can also modify large-scale recruitment of epileptic networks. To do this, we computed the mean number of propagation channels per IED during each arousal for all patients (*n* = 17). A propagation channel was defined as any SEEG channel in which an IED was recorded within a 120-ms window from the source IED.^11,30^ We then modelled the number of propagation channels using a GLMM, with condition (evoked arousal vs control) as the fixed effect of primary interest, adjusting for pre-stimulus number of propagation channels and stimulus intensity, and including a random intercept for patient ID. In contrast to the effects of evoked arousals on IED counts, propagation channel number did not differ significantly between evoked arousals and controls (GLMM, β-estimate ± SE: 0.212 ± 0.408 SE, IRR = 1.24, *P* = 0.604, **Fig. 3e**). Spontaneous arousals similarly showed no significant difference in number of propagation channels between arousals and controls (GLMM, β-estimate ± SE: 0.109 ± 0.317 SE, IRR = 1.12, *P* = 0.733, **Supplementary Fig. 6b**). Taken together, these results indicate that, while evoked arousals acutely increase interictal spiking by enhancing local cortical excitability, they do not adequately engage broader epileptic networks required to support widespread propagation of IEDs.

### Global interictal spiking is elevated during stimulation-induced sleep fragmentation

We then assessed whether the acute effects of evoked arousals on IEDs extended to a larger temporal scale. To do so, we compared global patient-level IED rates during NREM sleep between the first half of the sleep night, when the auditory stimulation protocol was delivered to induce sleep fragmentation, and the second half of the night, when auditory stimulation was withheld allowing recovery sleep. Normalised IED rates were significantly higher during stimulation-induced sleep fragmentation (0.80 ± 0.56 events/min (median ± IQR), range 0.32-1.50 events/min) compared to periods without auditory stimulation (0.72 ± 0.48 events/min, range 0.25-1.44 events/min) (Wilcoxon signed rank test, *p* = 0.017, Cliff’s *d* = 0.15, *n* = 17, **Supplementary Fig. 8a**). This pattern was confirmed using randomly sampled, arousal-free 3-s baseline epochs across the night: normalised IED counts were significantly higher during stimulation-induced sleep fragmentation than during periods without stimulation (GLMM, β-estimate ± SE: -0.112 ± 0.003, rate ratio = 0.89, *p* < 0.0001, **Supplementary Fig. 8b**), corresponding to a 10.6% reduction in IED count when auditory stimulation was stopped. These findings indicate that the broader disruption of sleep macrostructure caused by the auditory stimulation protocol is associated with elevated global IED activity over longer timescales beyond the immediate arousal window.

### Arousal-driven interictal spiking is dependent on anatomical region

Next, we asked whether arousal-driven increases in IEDs were dependent on anatomical region. To do this, we contrasted IEDs originating in mesiotemporal SEEG contacts (including the hippocampus, parahippocampal gyrus, entorhinal cortex, and amygdala) to those in the neocortex (**Fig. 4a,b**).^11^ This comparison allowed us to examine whether arousal-related IED increases differed between mesiotemporal and neocortical regions, which are implicated in epilepsy through distinct mechanisms,^41^ and differ in their functional connectivity with the thalamus, a key structure for sleep regulation.^42^ We modelled IED counts per channel-epoch using a GLMM, with condition (evoked arousal vs control), anatomical region (mesiotemporal vs neocortical), and their interaction as fixed effects, adjusting for pre-stimulus baseline IED count and stimulus intensity, and including random intercepts for SEEG contact nested within patient. This revealed a significant interaction between condition and anatomical region (GLMM, β-estimate ± SE: 0.737 ± 0.041, IRR = 2.09, *p* < 0.0001, **Fig. 4c**), indicating that the effect of evoked arousals on IED counts differed between mesiotemporal and neocortical regions. Post-hoc contrasts showed that, in patients with ≥3 mesiotemporal channels (*n* = 14), predicted IED count in mesiotemporal regions did not differ significantly between evoked arousals and controls (β-estimate = -0.087 ± 0.039, IRR = 0.92, *P* = 0.054, **Fig. 4c**). In contrast, neocortical IEDs (*n* = 17 patients) showed a significant 92% increase in predicted IED count during evoked arousals relative to controls (β-estimate = 0.650 ± 0.013, IRR = 1.92, *p* < 0.0001, **Fig. 4c**).

**Figure 4.**
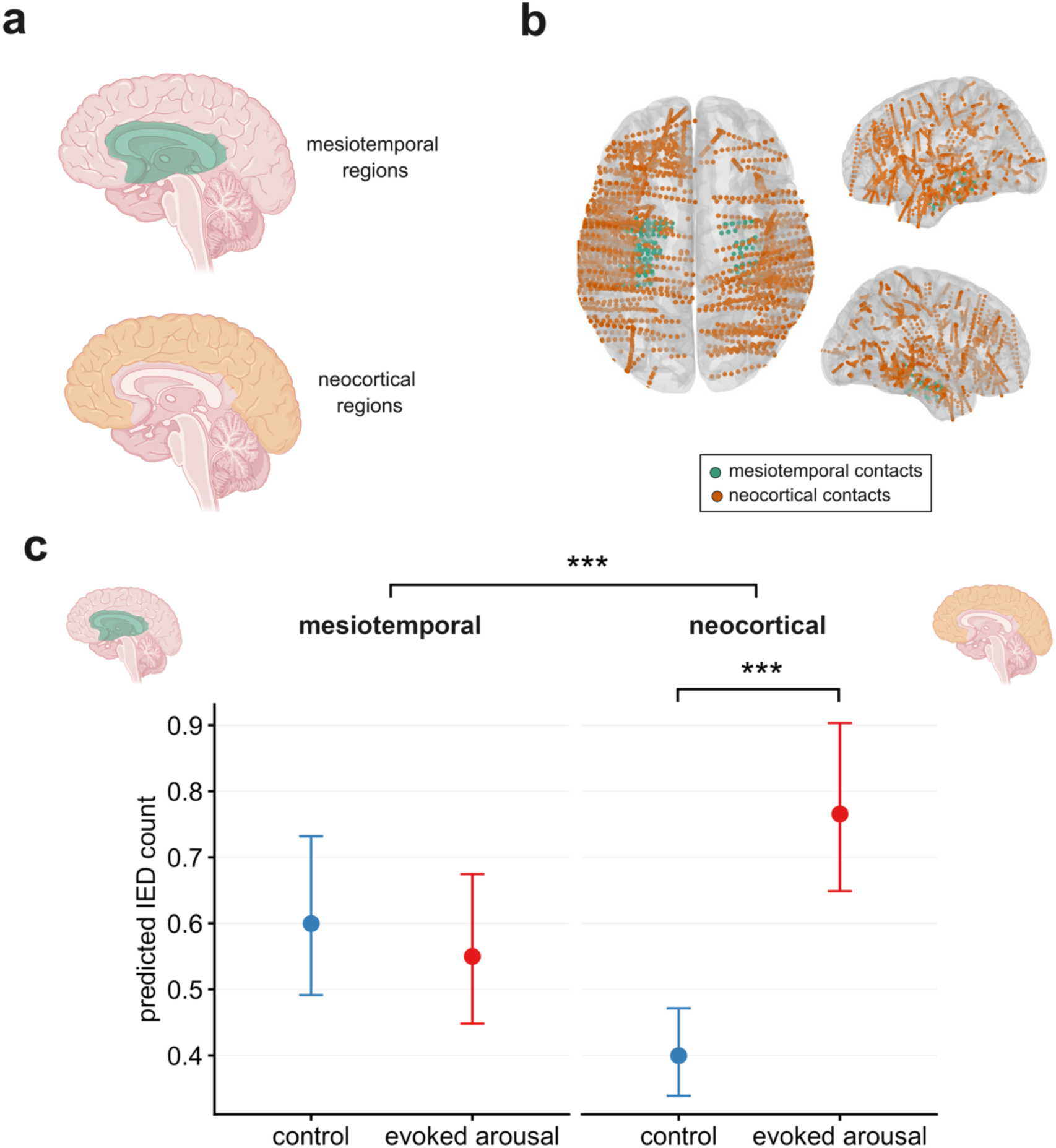
Arousal-related increases in interictal epileptiform discharges (IEDs) occur in neocortical but not mesiotemporal regions. **(a)** Schematic highlighting the two anatomical categories used for regional analysis: mesiotemporal (green) and neocortical (orange) regions were contrasted. **(b)** Spatial distribution of SEEG contacts across all patients (*n* = 17 patients), projected onto a standard brain template, showing mesiotemporal (green) and neocortical (orange) contact locations. **(c)** Model-predicted mean IED counts per channel-epoch by anatomical region (mesiotemporal vs neocortical) under control and evoked arousal conditions, adjusted for pre-stimulus IED count and stimulus intensity. The interaction between condition and anatomical region was significant (generalised linear mixed-effects model (GLMM), β-estimate ± standard error (SE): 0.737 ± 0.041, incidence rate ratio (IRR) = 2.09, *p* < 0.0001), indicating that the effect of evoked arousals on IED count differed between mesiotemporal and neocortical regions. In mesiotemporal regions (*n* = 14 patients), predicted IED count did not differ significantly between evoked arousals and controls (GLMM, β-estimate ± SE: -0.087 ± 0.039, IRR = 0.92, *P* = 0.054). In neocortical regions (*n* = 17 patients), predicted IED count was significantly higher during evoked arousals than controls (GLMM, β-estimate ± SE: 0.650 ± 0.013, IRR = 1.92, *p* < 0.0001), corresponding to a 92% increase. Error bars represent 95% confidence intervals. *** *p* < 0.001.

To assess whether this regional selectivity generalised beyond evoked arousals, we applied an equivalent GLMM framework to spontaneous arousals, which similarly revealed a significant interaction between condition and anatomical region (GLMM, β-estimate = 0.455 ± 0.030, IRR = 1.58, *p* < 0.0001, **Supplementary Fig. G**). However, in contrast to the evoked arousal findings, post-hoc contrasts showed a significant 27% decrease in predicted IED count in mesiotemporal regions during spontaneous arousals relative to controls (β-estimate = -0.320 ± 0.028, IRR = 0.73, *p* < 0.0001), alongside a significant 14% increase in neocortical regions (β-estimate = 0.135 ± 0.011, IRR = 1.14, *p* < 0.0001). We similarly examined regional selectivity for HFOs, and found a significant interaction between condition and anatomical region for evoked arousals (GLMM, β-estimate = 0.664 ± 0.050, IRR = 1.94, *p* < 0.0001, **Supplementary Fig. 10**), with no significant difference in mesiotemporal HFO count between evoked arousals and controls (β-estimate = -0.054 ± 0.048, IRR = 0.95, *P* = 0.511), but a significant 84% increase in neocortical HFO count (β-estimate = 0.610 ± 0.015, IRR = 1.84, *p* < 0.0001), mirroring the pattern observed for IEDs. Together, these findings show that arousal-related increases in interictal activity, including IEDs and HFOs, are preferentially expressed in the neocortex following evoked arousals, while mesiotemporal areas remain largely unaffected, highlighting regional differences in the sensitivity to transient sleep disruption. Notably, spontaneous arousals were instead associated with a significant decrease in mesiotemporal IED activity alongside a smaller yet still significant increase in the neocortex, suggesting that mesiotemporal and neocortical regions may respond differently depending on arousal type.

### Arousal-related interictal spiking is not modulated by the SOZ

We then investigated whether arousal-driven IED increases differed between SEEG contacts within the SOZ and those outside it (**Fig. 5a,b**). The SOZ was defined clinically based on the earliest electrographic changes at seizure onset.^43^ This comparison allowed us to determine whether the SOZ as a functional electrographic construct responds differently to arousals than surrounding areas, and whether it can modulate arousal-driven IEDs or remains resistant to these transient brain state changes. For patients who had a clear clinically identifiable SOZ in ≥3 SEEG channels (*n* = 12), we modelled IED counts per channel-epoch using a GLMM, with condition (evoked arousal vs control), SOZ status (outside vs inside SOZ), and their interaction as fixed effects, adjusting for pre-stimulus baseline IED count and stimulus intensity, and including random intercepts for SEEG contact nested within patient. This revealed no significant interaction between condition and SOZ status (GLMM, β-estimate ± SE: 0.005 ± 0.026, IRR = 1.00, *P* = 1.00, **Fig. 5**), indicating that the effect of evoked arousals on IED count did not differ significantly between SOZ and non-SOZ regions. Given the absence of a significant interaction, post-hoc contrasts were not performed separately for SOZ and non-SOZ regions.

**Figure 5.**
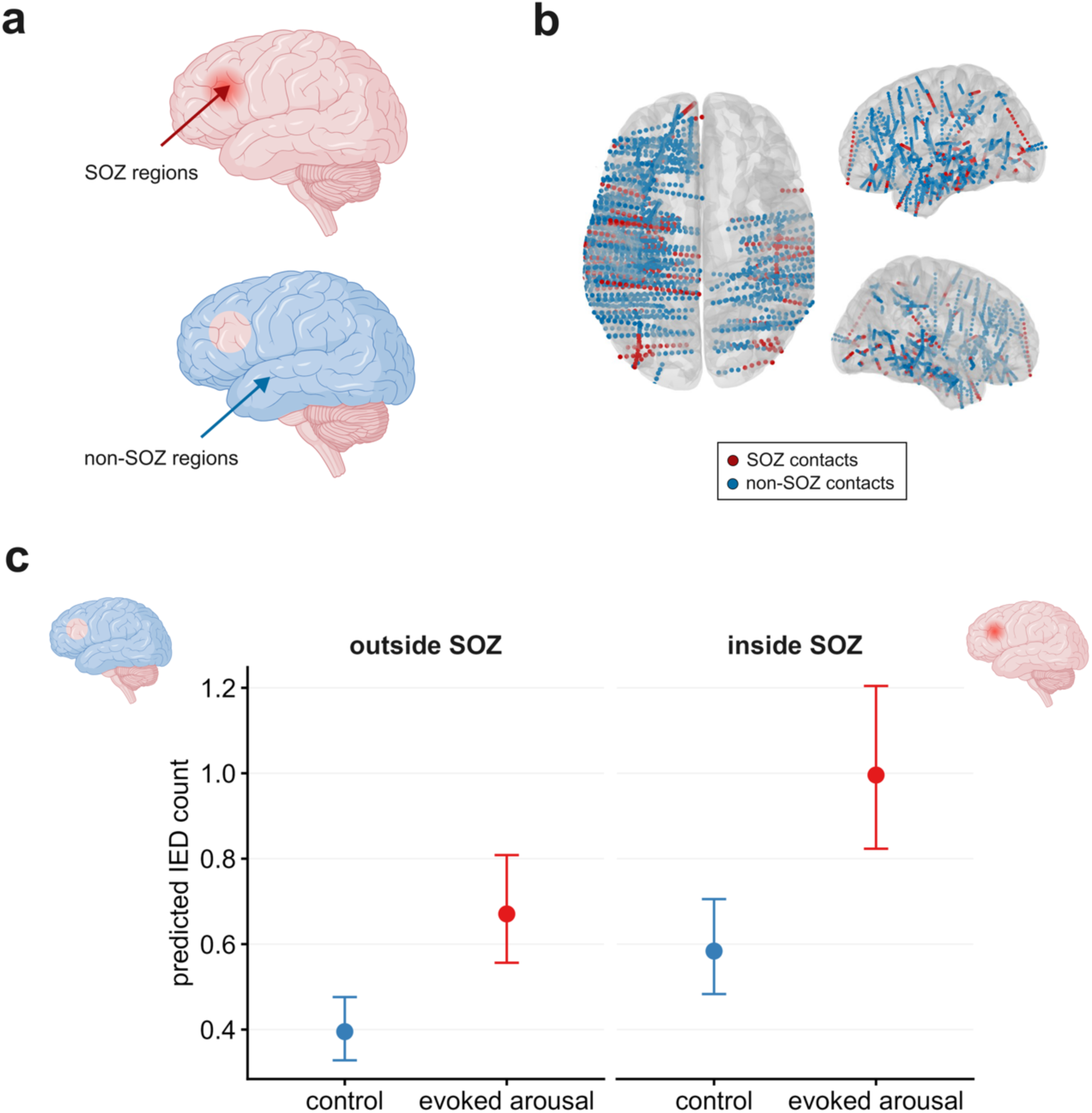
Arousal-related increases in interictal epileptiform discharges (IEDs) are not modulated by the seizure-onset zone (SOZ). **(a)** Schematic showing anatomical categorisation of SEEG contacts: IEDs originating in SOZ regions (red) and non-SOZ regions (blue) were contrasted. **(b)** Spatial distribution of SEEG contacts across all patients with an identifiable SOZ (*n* = 12), projected onto a standard brain template, showing SOZ (red) and non-SOZ (blue) contact locations. **(c)** Model-predicted mean IED counts per channel-epoch by SOZ status (outside vs inside SOZ) under control and evoked arousal conditions, adjusted for pre-stimulus IED count and stimulus intensity. The interaction between condition and SOZ status was not significant (generalised linear mixed-effects model (GLMM), β-estimate ± standard error (SE): 0.005 ± 0.026, incidence rate ratio (IRR) = 1.00, *P* = 1.00), indicating that the effect of evoked arousals on IED count did not differ between contacts inside and outside the SOZ. Error bars represent 95% confidence intervals.

Spontaneous arousals similarly revealed no significant interaction between condition and SOZ status (GLMM, β-estimate = -0.025 ± 0.021, IRR = 0.98, *P* = 0.940, **Supplementary Fig. 11**). We also observed a consistent pattern for HFOs, which again showed no significant interaction between condition and SOZ status for evoked arousals (GLMM, β-estimate = -0.056 ± 0.030, IRR = 0.95, *P* = 0.239, **Supplementary Fig. 12**). Together, these results indicate that arousal-related increases in interictal activity, including IEDs and HFOs, occur uniformly inside and outside the SOZ, suggesting that the SOZ does not modulate arousal-related brain state changes differently to surrounding tissue.

### Arousal-driven interictal spiking is dependent on sleep stage

We then asked whether the impact of arousals on IEDs depends on the sleep stage in which they occur. We contrasted arousals occurring during NREM stage 2 (N2) and stage 3 (N3) sleep (**Fig. 6a**), which represent distinct levels of neuronal synchrony and cortical excitability. While N2 is a lighter sleep stage characterised by sleep spindles and K-complexes, N3 (slow-wave sleep) is dominated by high-amplitude, low-frequency (0.5-2 Hz) delta activity and reduced cortical responsiveness.^13,44^ These neurophysiological differences may influence responses to evoked arousals. We modelled IED counts per channel-epoch using a GLMM, with condition (evoked arousal vs control), sleep stage (N2 vs N3), and their interaction as fixed effects, adjusting for pre-stimulus baseline IED count and stimulus intensity, and including random intercepts for SEEG contact nested within patient. This revealed a significant interaction between condition and sleep stage (GLMM, β-estimate ± SE: -0.271 ± 0.031, IRR = 0.76, *p* < 0.0001, **Fig. 6b**), indicating that the effect of evoked arousals on IED count differed between N2 and N3 sleep. Specifically, the arousal-related increase in IED count was approximately 24% smaller in N3 than in N2 sleep. Post-hoc contrasts showed that predicted IED count was significantly higher during evoked arousals relative to controls in both N2 sleep (*n* = 16 patients; β-estimate = 0.718 ± 0.017, IRR = 2.05, *p* < 0.0001, **Fig. 6b**) and N3 sleep (*n* = 13 patients; β-estimate = 0.447 ± 0.026, IRR = 1.56, *p* < 0.0001, **Fig. 6b**), though this arousal-related increase was significantly larger in N2 than in N3 sleep.

**Figure 6.**
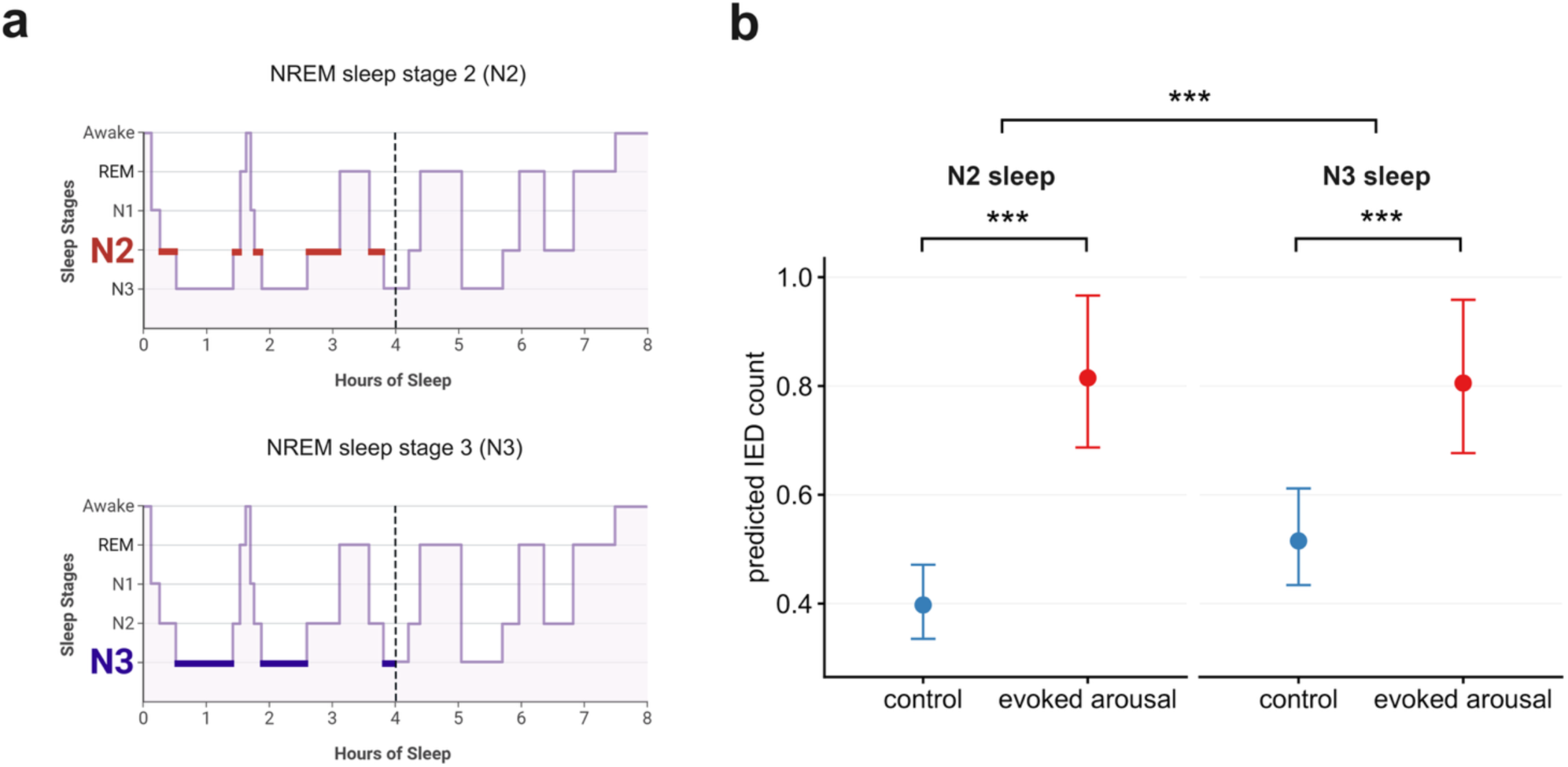
Arousal-related increases in interictal epileptiform discharges (IEDs) are sleep-stage dependent. **(a)** Schematic hypnograms with NREM stage 2 (N2) and stage 3 (N3) sleep highlighted during the first half of the night, defined as the first 4 hours following each patient’s habitual bedtime, when the auditory stimulation protocol was delivered. Evoked arousals occurring during N2 and N3 sleep were contrasted to assess sleep-stage effects on arousal-driven interictal epileptiform discharges (IEDs). **(b)** Model-predicted mean IED counts per channel-epoch by sleep stage (N2 vs N3 sleep) under control and evoked arousal conditions, adjusted for pre-stimulus IED count and stimulus intensity. The interaction between condition and sleep stage was significant (generalised linear mixed-effects model (GLMM), β-estimate ± standard error (SE): -0.271 ± 0.031, incidence rate ratio (IRR) = 0.76, *p* < 0.0001), indicating that the effect of evoked arousals on IED count differed between N2 and N3 sleep, corresponding to a 24% smaller increase in N3 compared to N2. In N2 sleep (*n* = 16 patients), predicted IED count was significantly higher during evoked arousals than controls (GLMM, β-estimate ± SE: 0.718 ± 0.017, IRR = 2.05, *p* < 0.0001). In N3 sleep (*n* = 13 patients), predicted IED count was also significantly higher during evoked arousals than controls though with a smaller magnitude than in N2 (GLMM, β-estimate ± SE: 0.447 ± 0.026, IRR = 1.56, *p* < 0.0001). Error bars represent 95% confidence intervals. *** *p* < 0.001.

We then examined whether this sleep stage-dependent pattern generalised to spontaneous arousals. Applying an equivalent GLMM framework to spontaneous arousals, we found a significant interaction between condition and sleep stage (GLMM, β-estimate = 0.089 ± 0.029, IRR = 1.09, *P* = 0.0075, **Supplementary Fig. 13**). Post-hoc contrasts showed that predicted IED count was significantly higher during spontaneous arousals relative to controls in both N2 sleep (β-estimate = 0.143 ± 0.015, IRR = 1.15, *p* < 0.0001) and N3 sleep (β-estimate = 0.232 ± 0.025, IRR = 1.26, *p* < 0.0001). Notably, the spontaneous arousal-related increase in IED count was 10% larger in N3 than in N2 sleep, the reverse of that observed for evoked arousals.

For HFOs, we observed a consistent pattern to the evoked-arousal IED findings, with a significant interaction between condition and sleep stage (GLMM, β-estimate = -0.275 ± 0.036, IRR = 0.76, *p* < 0.0001, **Supplementary Fig. 14**). Post-hoc contrasts showed that predicted HFO count was significantly higher during evoked arousals relative to controls in both N2 sleep (β-estimate = 0.758 ± 0.020, IRR = 2.13, *p* < 0.0001) and N3 sleep (β-estimate = 0.483 ± 0.030, IRR = 1.62, *p* < 0.0001), with the arousal-related increase approximately 24% smaller in N3 than in N2, mirroring the pattern observed for IEDs during evoked arousals.

Together, these results indicate that arousals increase epileptic activity across both N2 and N3 sleep, but that the relative magnitude of this effect is sleep-stage dependent. For both IEDs and HFOs during evoked arousals, this effect was significantly greater in N2 than N3 sleep, consistent with N2 representing a state of relatively higher cortical excitability and network susceptibility to transient sleep instability. Interestingly, this pattern was reversed for spontaneous arousals, which were associated with a significantly larger increase in IED count during N3 than N2 sleep, suggesting that the mechanisms underlying evoked and spontaneous arousal-related increases in interictal activity may differ in their sensitivity to sleep stage.

### Influence of stimulus response type and pre-stimulus delta activity on interictal spiking

We then tested whether the facilitation of IEDs during arousals was linked to the arousal itself or the accompanying slow-wave response, and whether pre-stimulus delta activity conditioned the electrophysiological and epileptiform response to stimulation. This addressed whether the observed IED increases are specific to conventional fast arousals, or reflect a broader form of stimulus-induced instability in slow oscillations that also occurs when a measurable slow-wave response is elicited without meeting formal arousal criteria.^13,45^ It also allowed us to examine whether pre-stimulus delta activity, as a marker of background instability, influenced the likelihood of an arousal response and any subsequent increase in IEDs.

To address these questions, we fitted two further models. First, a GLMM was used to test whether pre-stimulus spectral power ratio (δ/[θ+α+β]; [0.5–4 Hz]/[4.5–30 Hz]) predicted the probability of an arousal response relative to an isolated slow-wave response, with pre-stimulus spectral power ratio as the fixed effect of interest, adjusting for stimulus intensity and pre-stimulus baseline IED count, and including random intercepts for SEEG contact nested within patient. As stimuli with no detectable EEG response occurred in only 4 patients, this category was excluded, allowing all 17 patients to be retained in a comparison of two response types: isolated slow wave and arousal with slow-wave activity. The distributions of the pre-stimulus spectral power ratio were broadly similar between the two response types (**Fig 7a**), and this was confirmed statistically: pre-stimulus spectral power ratio was not associated with the likelihood of an arousal response relative to an isolated slow-wave response (GLMM, β-estimate ± SE: 0.032 ± 0.014, *P* = 0.052, **Fig. 7b**).

**Figure 7.**
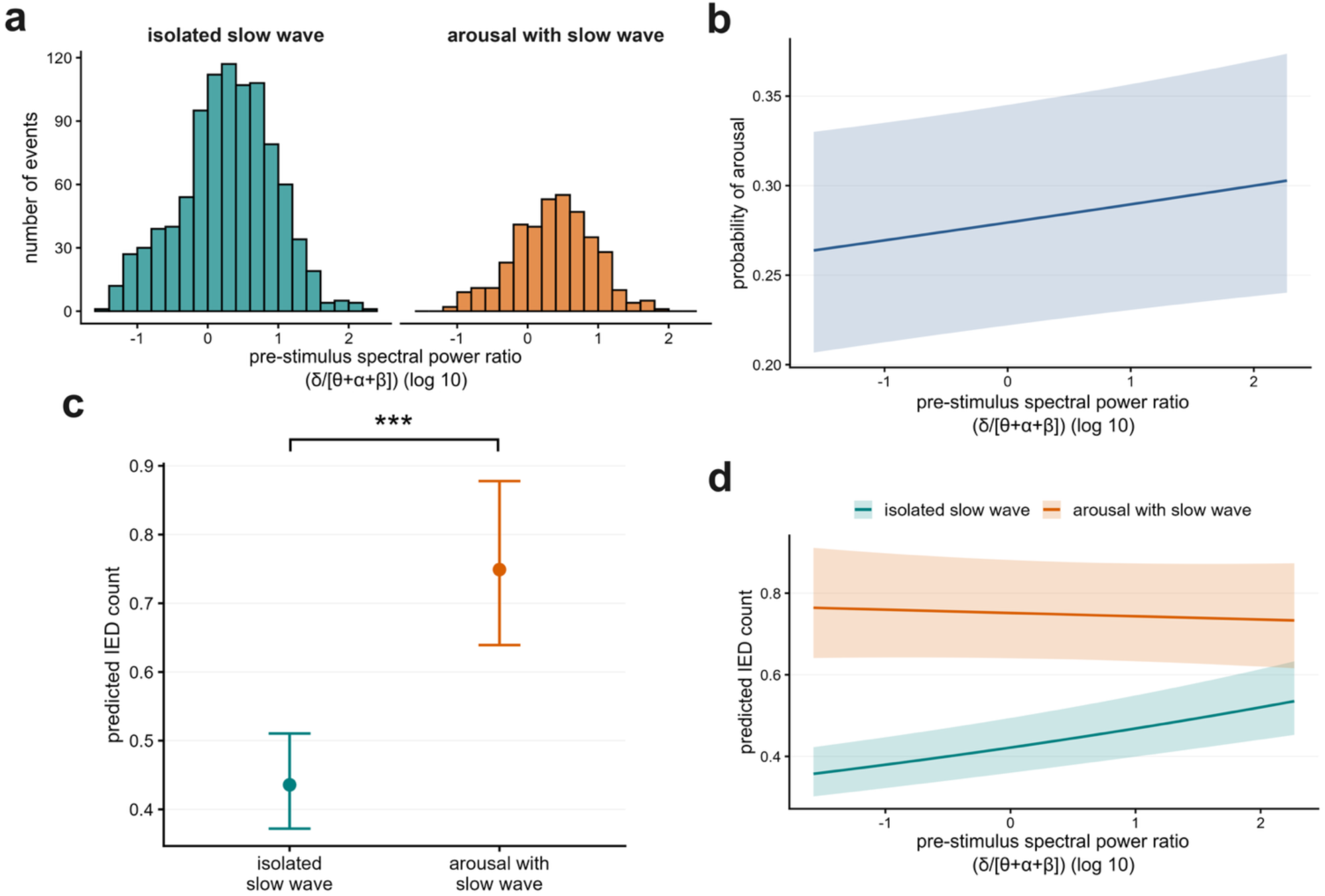
Interictal epileptiform discharges (IEDs) are primarily driven by arousal responses rather than isolated slow waves, with pre-stimulus delta activity exerting a condition-specific modulatory effect. **(a)** Histograms displaying the distribution of the pre-stimulus spectral power ratio (δ/[θ+α+β] ([0.5–4 Hz]/[4.5–30 Hz], log10-transformed) preceding auditory stimuli that resulted in isolated slow waves and arousals with slow waves. **(b)** Model-predicted probability of an arousal response to stimulation as a function of pre-stimulus spectral power ratio, adjusted for stimulus intensity and pre-stimulus baseline IED count. Pre-stimulus spectral power ratio was not associated with the likelihood of an arousal response relative to an isolated slow-wave response (generalised linear mixed-effects model (GLMM), β-estimate ± standard error (SE): 0.032 ± 0.014, *P* = 0.052). **(c)** Model-predicted mean IED counts for isolated slow waves and arousals with slow waves. Arousals with slow waves were associated with significantly higher IED counts compared to isolated slow waves (GLMM, β-estimate ± SE: 0.542 ± 0.012, incidence rate ratio (IRR) = 1.72, *p* < 0.0001), corresponding to a 72% increase in IED count. **(d)** The interaction between response type and pre-stimulus spectral power ratio was also significant (GLMM, β-estimate ± SE: -0.073 ± 0.013, IRR = 0.93, *p* < 0.0001). Post-hoc simple slopes analysis revealed that a higher pre-stimulus spectral power ratio significantly increased IED counts during isolated slow waves (GLMM, β-estimate ± SE = 0.066 ± 0.009, IRR = 1.07, *p* < 0.0001), corresponding to a 7% increase in IED count per 1.0 increase in pre-stimulus spectral power ratio. In contrast, pre-stimulus spectral power ratio did not influence IED counts once an arousal was triggered (GLMM, β-estimate ± SE = -0.007 ± 0.012, IRR = 0.99, *P* = 1). Error bars and shaded areas represent 95% confidence intervals. *** *p* < 0.001.

We then used a GLMM to evaluate the effect of stimulus response type on IED count, and to test whether pre-stimulus spectral power ratio modulated this relationship. IED count was modelled with response type (isolated slow wave vs arousal with slow wave), pre-stimulus spectral power ratio, and their interaction as fixed effects, adjusting for pre-stimulus baseline IED count and stimulus intensity, and including random intercepts for SEEG contact nested within patient. Arousals with slow-wave activity were associated with significantly higher IED counts than isolated slow waves (GLMM, β-estimate ± SE: 0.542 ± 0.012, IRR = 1.72, *p* < 0.0001, **Fig. 7c**), corresponding to a 72% increase in IED count. The interaction between response type and pre-stimulus spectral power ratio was also significant (GLMM, β-estimate ± SE: -0.073 ± 0.013, IRR = 0.93, *p* < 0.0001, **Fig. 7d**), indicating that the relationship between pre-stimulus delta activity and IED count depended on the type of response elicited. Post-hoc simple slopes analysis was performed to characterise this relationship separately within each response type. Following an isolated slow-wave response, a higher pre-stimulus spectral power ratio was associated with a significantly greater IED count (β-estimate = 0.066 ± 0.009, IRR = 1.07, *p* < 0.0001, **Fig. 7d**), corresponding to a 7% increase in IED count per 1.0 increase in pre-stimulus spectral power ratio, indicating that a more delta-dominant pre-stimulus state increased epileptiform activity in the absence of an arousal. In contrast, following an arousal, pre-stimulus spectral power ratio was not significantly associated with IED count (β-estimate = -0.007 ± 0.012, IRR = 0.99, *P* = 1, **Fig. 7d**), indicating that once an arousal occurred, IED count was largely independent of the preceding delta state. Together, these findings suggest that baseline delta activity acts as a condition-specific modulator of epileptiform activity: it facilitates IED increases following isolated slow waves, but has no additional influence once an arousal response is triggered.

Given that pre-stimulus delta activity showed a condition-specific relationship with IED count, we next tested the possibility that it also represented a direct confounding pathway between response type and IED count, using a change-in-estimate sensitivity analysis. The inclusion of the pre-stimulus delta activity did not alter the estimated effect of stimulus response type on IED count when comparing the unadjusted model (GLMM, β-estimate ± SE: 0.538 ± 0.012, IRR = 1.71, *p* < 0.0001) with the model adjusted for pre-stimulus spectral power ratio (GLMM, β-estimate ± SE: 0.537 ± 0.012, IRR = 1.71, *p* < 0.0001). Furthermore, the pre-stimulus spectral power ratio was not a significant independent predictor of IED count in the adjusted model (β-estimate = 0.009 ± 0.005, IRR = 1.01, *P* = 0.081). This demonstrates that, under a linear assumption, pre-stimulus delta activity does not act as a primary confounder in the relationship between response type and IED count, supporting response type rather than background brain state as the primary driver of epileptiform activity following stimulation.

To further address the possibility that stimulus intensity may itself be driving IED counts, we examined whether stimulus intensity was associated with a change in IED count during control epochs, where auditory stimuli did not result in a detectable arousal response. During control epochs, stimulus intensity was associated with a small but significant increase in IED count (GLMM, β-estimate ± SE: 0.0056 ± 0.0005, IRR = 1.006, *p* < 0.0001, **Supplementary Fig. 15a**), corresponding to a 0.6% increase in IEDs per 1 dB increase in stimulus intensity. However, the interaction between event type (evoked arousal vs control) and stimulus intensity was not significant (GLMM, β-estimate ± SE: - 0.0004 ± 0.0006, IRR = 1.00, *P* = 0.514, **Supplementary Fig. 15b**), indicating that the larger IED increases observed during evoked arousals represent a distinct state change that occurs independently of stimulus intensity. Moreover, stimulus intensity was not significantly associated with change in thalamic spectral power ratio in control epochs, as measured in patients who had SEEG electrodes implanted within the thalamus (*n* = 4 patients; GLMM, β-estimate ± SE: -0.008 ± 0.009, *P* = 0.382, **Supplementary Fig. 15c**), suggesting that stimulus intensity did not drive detectable shifts in thalamic spectral content in the absence of an arousal response. Together, these findings indicate that while control epochs were associated with a small, intensity-dependent increase in IED count – potentially reflecting sub-threshold arousal-like responses or direct sensory activation not evident on the PSG – this effect did not scale with, or account for, the substantially larger and distinct increase in IED activity observed during evoked arousals.

### Temporal dynamics of arousal-driven interictal spiking

Having established that interictal spiking increases during transient arousals, we next examined the temporal dynamics of this effect. We asked whether the number and timing of IEDs vary as a function of arousal duration and whether IED increases are time-locked to arousal onset or distributed across the full arousal. This analysis was designed to characterise how interictal activity evolves over time relative to arousal state transitions, thereby providing insights into the dynamic interplay between cortical activation and pathological excitability. To visualise these dynamics, we quantified the temporal distribution of IEDs relative to arousal and stimulus onset across patients (*n* = 17), with IED counts normalised to each patient’s maximum value and aggregated in 500-ms bins (**Fig. 8a-c**). IED counts increased significantly immediately following arousal onset, compared to the pre-arousal baseline, and remained elevated for 4.5 s of the post-arousal period (one-tailed Mann-Whitney tests, *p* < 0.05, **Fig. 8a**), peaking at 0.5-1 s after arousal onset. Patient-level distributions of normalised IED count aligned to arousal onset similarly showed a clear tendency for IEDs to cluster early in the arousal period, suggesting a rapid rise in cortical excitability immediately following arousal onset which stabilises over time (**Supplementary Fig. 16**). When aligned to auditory stimuli onset, IED counts again increased significantly during evoked arousals, peaking around 2-2.5 s after stimulus onset (one-tailed Mann–Whitney tests, *p* < 0.05, **Fig. 8b**). The proportion of active arousals over time closely mirrored this pattern, highlighting tight temporal coupling between arousal activation and IED increases. In contrast, no significant modulation of IED activity was observed during control non-arousing auditory stimuli (one-tailed Mann–Whitney tests, *p* > 0.05, **Fig. 8c**), confirming that the observed increase was specific to arousal-related cortical activation rather than the auditory stimulus itself.

**Figure 8.**
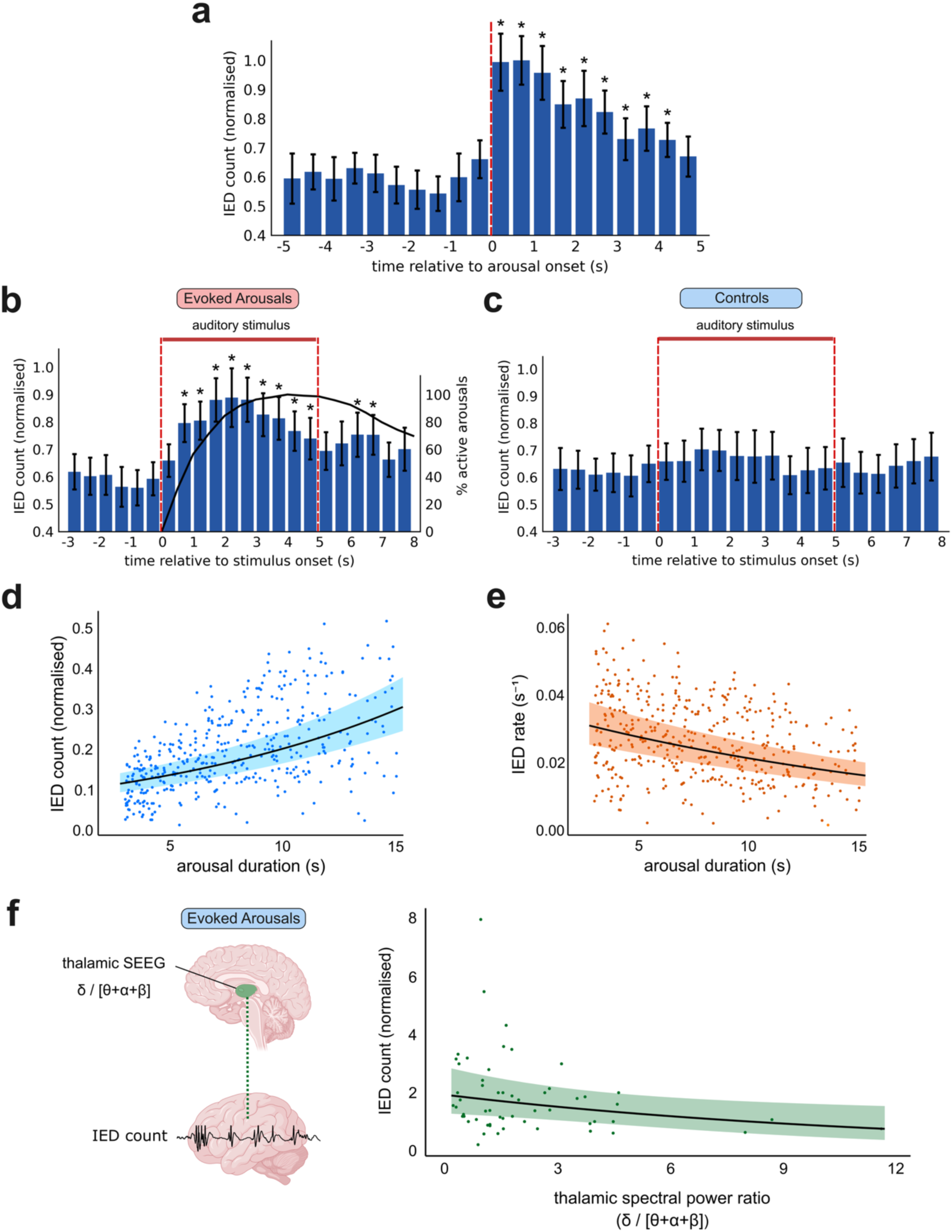
Temporal dynamics of interictal epileptiform discharges (IEDs) during evoked arousals. (a-c) Group-level histograms displaying the temporal distribution of IEDs relative to arousal and auditory stimulus onset, normalised to each patient’s maximum value (bin width: 500 ms) and averaged across all patients, with group data presented as mean ± standard error. IED counts during the pre-onset baseline period were compared to that during each post-onset bin using one-tailed Mann-Whitney tests (*n* = 17 patients, significant bins (*p* < 0.05) indicated by asterisks). **(a)** IED counts increased significantly following the onset of evoked arousals (red dashed line), peaking at 0.5-1 s after arousal onset. **(b)** Aligned to the onset of auditory stimuli, IED counts again increased significantly during evoked arousals, peaking at 2-2.5 s after stimulus onset. The overlaid black curve (right axis) shows the percentage of active arousals over time. **(c)** During control (non-arousing) auditory stimuli, no significant modulation of IED count was observed following stimulus onset, indicating that the IED increase observed in (b) was specific to evoked arousals. **(d)** IED count, normalised by number of SEEG contacts in each patient, increased significantly with arousal duration (generalised linear mixed-effects model (GLMM), β-estimate ± standard error (SE): 0.078 ± 0.006 per second, *p* < 0.0001), confirming that longer arousals are associated with higher IED count. **(e)** In contrast, IED rate decreased significantly with arousal duration (GLMM, β-estimate ± SE: –0.051 ± 0.006 per second, *p* < 0.0001), suggesting that although longer arousals contain more IEDs, they occur at a progressively slower rate over the course of the arousal. (**f**) Thalamic spectral activity is associated with arousal-related IED count. A subset analysis in four patients with thalamic SEEG electrodes (*n* = 56 arousals during N2 sleep) showed that a higher thalamic spectral power ratio (δ/[θ+α+β] ([0.5–4 Hz]/[4.5–30 Hz])) was associated with a lower normalised IED count (GLMM, β-estimate ± SE: –0.081 ± 0.035 per 1.0 increase in spectral power ratio, *P* = 0.022), indicating that a shift toward faster thalamic rhythms during arousals is linked to greater interictal spiking. Each dot in panels (**d-f**) represents a single arousal event, pooled across patients, and shaded regions represent the 95% confidence intervals around the model fit.

To assess the relationship between arousal duration and interictal spiking, we pooled arousals and associated IEDs across patients (*n* = 17) and performed generalised linear mixed-effects modelling (GLMM) with arousal duration as the independent variable and normalised IED count or IED rate as the dependent variable. IED count increased significantly with arousal duration (GLMM, β-estimate ± SE: IED count increased by 0.078 ± 0.006 with each 1-s increase in arousal duration, *p* < 0.0001, **Fig. 8b**), indicating that longer arousals are associated with more interictal activity overall. Interestingly, IED rate across the full arousal decreased significantly with arousal duration (GLMM, β-estimate ± SE: IED rate decreased by 0.051 ± 0.006 s^-1^ with each 1-s increase in arousal duration, *p* < 0.0001, **Fig. 8c**). This suggests that while longer arousals contain more IEDs, they occur at a progressively slower rate as the arousal progresses. Together, these findings indicate that arousal onset is associated with a transient surge in epileptic activity that diminishes as the arousal stabilises, reflecting that arousal-induced interictal spiking is driven by dynamic fluctuations in cortical excitability.

Finally, we examined whether arousal-driven increases in interictal activity can be explained by thalamocortical activation, by performing a subset analysis in four patients who had SEEG electrodes implanted within the thalamus. We applied a GLMM to test whether thalamic spectral power was associated with IED counts during N2 arousals (*n* = 56), where a significantly larger increase in IED count had been observed. Normalised IED count across all SEEG channels was modelled as a function of the thalamic spectral power ratio, δ/(θ+α+β) ([0.5–4 Hz]/[4.5–30 Hz]), computed over the 3-s arousal epoch normalised to the pre-stimulus baseline. We found a significant negative relationship between thalamic spectral power ratio and IED count (GLMM, β-estimate ± SE: IED count decreased by 0.081 ± 0.035 with each 1.0 increase in thalamic spectral power ratio, *P* = 0.022, **Fig. 8f**). Therefore, arousals accompanied by lower relative delta activity – reflecting a shift toward faster thalamic rhythms – were associated with higher IED counts. These results indicate that arousal-driven interictal spiking is linked to transient thalamic activation, whereby the emergence of faster thalamic oscillations may facilitate cortical excitability and the generation of IEDs. Collectively, these findings provide a mechanistic bridge between thalamocortical signatures of arousal and the expression of arousal-related interictal activity.

## Discussion

In summary, our findings reveal that sleep arousals are a critical and dynamic modulator of interictal activity. Using combined SEEG-PSG recordings and controlled auditory stimulation to manipulate sleep stability and microstructure, we demonstrate that evoked arousals acutely increase pathological discharges in a network-specific manner. Although complex bidirectional links between epilepsy and sleep are recognised and prior work has demonstrated close associations between sleep instability and epileptic activity,^1,3,8,19^ the causal impact of sleep instability on epileptic activity had not been established. By experimentally evoking arousals, our study provides direct perturbational evidence that transient increases in sleep instability cause an immediate increase in IEDs. This effect generalises to HFOs and spontaneous arousals not induced by auditory stimulation. Notably, while arousals transiently enhance IED occurrence, they do not alter their spatial propagation, suggesting that sleep fragmentation modulates epileptic activity without involving widespread network recruitment. We propose that this reflects a transient elevation in local cortical excitability that facilitates the emergence of epileptic activity,^46,47^ supported by our finding that arousal-related IED increases are specific to neocortical regions, and absent in mesiotemporal areas, reflecting a region-dependent effect. We further show that arousal-driven IED increases do not differ between the SOZ and surrounding regions, suggesting that the SOZ is not preferentially modulated by transient fluctuations in cortical excitability associated with sleep instability.^30^ Moreover, arousal-driven increases in IEDs show sleep-stage dependence, occurring in both N2 and N3 sleep but with a larger effect in N2, consistent with stronger thalamocortical synchronisation in N2 sleep which may promote epileptic activity.^48,49^ Results from a subset of patients with thalamic SEEG electrodes indicate that faster thalamic activity during N2 arousals is associated with higher IED counts, consistent with a link between transient thalamocortical activation and neocortical interictal spiking in this sleep stage. Finally, we show that while background delta activity facilitates IEDs during isolated slow-wave responses, this modulatory effect is overridden once an arousal occurs, highlighting sleep arousals as the dominant drivers of interictal spiking during fragmented sleep.

The immediate increase in IED count during evoked arousals likely reflects a shift in cortical excitability associated with abrupt sleep transitions, consistent with prior work linking sleep instability in the context of the CAP and arousal-related dynamics to increased epileptic activity.^3,14–16^ A consistent increase was observed for HFOs, and for spontaneous arousals, indicating that this reflects a broader influence of sleep instability on epileptiform activity rather than one specific to auditory-evoked events or to IEDs alone. Sleep arousals are defined by a temporary shift towards higher EEG frequencies, representing an intermediate state between stable sleep and full wakefulness.^13,50^ This may temporarily destabilise the excitatory-inhibitory balance within the cortical network, and promote a hyperexcitable state conducive to IED generation.^47,51^ Given the brief duration of arousals, the associated increase in cortical excitability may be too short-lived to give rise to seizures, potentially explaining their absence during arousals, while still driving IED increases. This is further supported by our finding that IED rates peak immediately following arousal onset and decline as the arousal progresses, suggesting that interictal spiking is temporally coupled to the initial unstable transition into the arousal state. Together, these findings support the view that transient increases in sleep instability – particularly during arousal transitions – can modulate epileptic activity on rapid timescales, extending prior observational and mechanistic work by providing direct perturbational evidence. These results highlight how sleep can serve as a powerful tool to study the dynamic regulation of epileptic activity.

Contrary to our hypothesis, evoked arousals did not influence the spatial propagation of IEDs, despite consistently increasing their occurrence; the same was true for spontaneous arousals. These differing effects suggest that arousals primarily modulate the local generation of interictal activity without recruiting broader epileptic networks that would facilitate its spread.^52,53^ Therefore, transient shifts in cortical excitability during arousals may act as a trigger to initiate IEDs in local networks, but do not reach the threshold required for widespread network activation.^54^ These findings reinforce the idea that IED generation and propagation are governed by distinct mechanisms; while IED generation may be facilitated by rapid arousal-related fluctuations in local cortical excitability, their propagation requires globally coordinated activation across brain regions mediated by larger-scale network dynamics.

Regional differences in arousal-driven IED increases can be explained by varying thalamocortical connectivity across brain regions. The neocortex, which receives dense thalamic projections and exhibits strong functional connectivity with arousal-regulating thalamic nuclei,^39,55,56^ showed IED increases during evoked arousals, whereas mesiotemporal regions showed no significant effect, possibly due to their comparatively limited connectivity to the thalamus.^56,57^ Interestingly, while spontaneous arousals were also accompanied by a significant increase in neocortical IEDs, they showed an IED decrease in mesiotemporal regions. Our findings align with previous work demonstrating a neocortical IED increase and mesiotemporal IED decrease during spontaneous arousals.^19^ The suppression of mesiotemporal IEDs during spontaneous, but not evoked, arousals may reflect differing combinations of central and peripheral components between arousal types.^8^ Auditory evoked arousals, for instance, may engage concurrent sensory processing pathways that counteract the desynchronisation of mesiotemporal network activity observed in endogenously generated arousals, thereby reducing IED suppression.^8,58^

The regional specificity between neocortical and mesiotemporal structures is further supported by analysis in a subset of patients with thalamic SEEG electrodes, which revealed that arousals accompanied by faster thalamic activity were associated with higher IED counts. These results suggest that transient thalamocortical activation during arousals may preferentially drive interictal spiking in the neocortex, providing a mechanistic explanation for the neocortical IED increase. While mesiotemporal structures including the hippocampus are strongly connected to the anterior thalamic nuclei as part of the Papez circuit,^59^ this pathway is typically implicated in memory and limbic function rather than in global arousal and state regulation.^60^ Neocortical activation during arousal, by contrast, is supported by thalamic systems strongly involved in the regulation of cortical state, including the midline and intralaminar nuclei.^61^ It is therefore possible that thalamic activity accompanying arousals engages these functionally distinct pathways to differing degrees, contributing to the regional differences in IED responses observed here. Future work incorporating larger thalamic samples with more detailed anatomical localisation of thalamic nuclei and their projection targets would help to further refine these mechanistic interpretations. Additionally, variations in local arousal patterns such as the presence of slow-wave activity, which is associated with IEDs, may further contribute to this regional specificity.^19^

The comparable arousal-driven IED increases within and outside the SOZ suggest that the transient excitability changes associated with sleep arousals are not selectively gated by pathological network alterations within the SOZ. This finding indicates that arousal-related modulation of interictal activity may reflect a global network phenomenon rather than being specifically amplified within seizure foci. This is consistent with previous studies showing that regions both within and outside the SOZ remaining dynamically coupled to changes in cortical excitability across vigilance states.^11,62^

Arousal-driven interictal spiking also showed sleep-stage specificity, with a significantly larger effect in N2 than N3 sleep, aligning with previous work that showed a stronger arousal-IED association in N2 and evidence that epileptic activity occurs more frequently during lighter, less stable N2 sleep, whereas it is reduced in deeper, more stable N3 sleep.^9,19–21^ N2 sleep is characterised by greater sleep instability, including more frequent cyclic alternating pattern (CAP)-related fluctuations and arousals, whereas N3 reflects a more stable form of synchronised sleep.^8,66,67^ In this context, the preferential increase in IEDs during N2 may reflect not only differences in network organisation but also the intrinsically higher instability of this sleep stage, which may facilitate transient increases in cortical excitability. This is further supported by findings that the temporal and spatial instability of delta activity, as opposed to just slow-wave power alone, are associated with increased epileptic activity.^20,21^ The difference between N2 and N3 may also be rooted in the distinct thalamocortical dynamics underlying these sleep stages. As a central regulator of sleep, the thalamus plays a crucial role in coordinating cortical excitability and responsiveness.^42,48^ N2 sleep is associated with stronger thalamocortical synchronisation, which may enhance the propagation of arousal-related excitatory activity and promote IED generation.^48,68^ In contrast, N3 sleep is marked by reduced thalamic engagement and more prominent cortico-cortical connectivity, potentially limiting the cortical network’s responsiveness to arousals.^48^ Notably, spontaneous arousals showed a significantly larger IED increase in N3 than N2, despite significant increases in both stages relative to control epochs, suggesting that auditory evoked and endogenously generated arousals may differ in their sensitivity to the underlying sleep state.^8^

Building on the link between delta activity and epileptiform activity,^20,21^ we asked whether IED facilitation reflects the arousal response itself or the accompanying slow-wave activity, and whether pre-stimulus delta state further modulates this relationship. Full cortical arousals were associated with significantly higher IED counts than isolated slow-wave responses, indicating that the additional cortical activation accompanying a full arousal, beyond slow-wave activity alone, is an important component of the observed acute increase in epileptiform activity. For isolated slow-wave responses, a more delta-dominant pre-stimulus state predicted greater IED counts, consistent with prior evidence linking delta activity to epileptiform discharges.^20,21,45^ Once an arousal occurred, however, IED count was no longer associated with the pre-stimulus delta state, suggesting that arousals exert a sufficiently strong influence on cortical excitability to override any additional contribution of background delta instability.^8^ Together, these findings point to two partially dissociable influences on epileptiform activity: background delta activity, most influential in the absence of a stronger perturbation, and arousal-related cortical activation, which appears to become the dominant influence once an arousal is triggered. This is consistent with the CAP framework and broader view that arousals exist on a continuum of intensity and EEG expression,^8,12,16^ and highlights the need for future work to better characterise arousal intensity and its physiological and pathological correlates.

While stimulus intensity had a small dose-dependent influence on IED count during control epochs, the larger IED increase observed during arousals occurred independently of stimulus intensity. Moreover, the absence of a corresponding change in thalamic spectral content alongside this stimulus-dependent increase suggests that stimulus intensity may modulate epileptiform activity through auditory sensory processing and subthreshold activation of ascending brainstem networks,^69,70^ rather than through thalamocortical modulation associated with evoked arousals.

Our findings advance understanding of sleep-epilepsy interactions by demonstrating that arousals act as acute, dynamic perturbations that destabilise cortical network excitability and drive interictal activity across both short and long timescales. Despite receiving less clinical attention than seizures, increasing evidence suggests that IEDs can disrupt memory consolidation, impair cognitive performance, and destabilise epileptic networks.^32–36^ Memory complaints are common even in patients without frequent seizures,^71–73^ and may reflect the cumulative impact of interictal activity during sleep. This is particularly important given that sleep plays a critical role in supporting the long-term consolidation of various forms of memory, including declarative and procedural memories.^37,38,74,75^ Indeed, elevated IEDs during NREM sleep have been proposed as a mechanistic contributor to accelerated long-term forgetting in patients with epilepsy,^76^ and may act as pro-epileptogenic events that lower the threshold for subsequent seizures.^77–79^ Nocturnal seizures, in turn, are linked to increased risk of SUDEP,^4^ highlighting the need to better understand and mitigate the upstream mechanisms that modulate interictal activity during sleep. Our results demonstrate that arousals act as a modifiable trigger for transient IED increases, with effects most prominent in neocortical regions and in N2 sleep, which has important implications for the clinical management of epilepsy. Reducing this arousal-linked interictal burden could be accomplished through several avenues. First, systematic screening and treatment of comorbid sleep fragmentation triggers, such as obstructive sleep apnoea,^80^ could directly eliminate a major source of arousal-induced spike surges. Secondly, targeted pharmacotherapy could prioritise sleep-consolidating medications or specific antiseizure medications that promote sleep continuity without disrupting critical NREM architecture.^81,82^ Finally, emerging technologies, such as closed-loop neuromodulation, could be adapted for real-time detection and suppression of NREM network instability. Notably, most sleep-focused interventions target slow-wave sleep (N3);^83–85^ however, our findings suggest that stabilising N2 sleep is also important for reducing IED burden and improving clinical outcomes in epilepsy. Increasing evidence shows that interictal activity is also implicated in Alzheimer’s disease and related dementias, highlighting pathophysiological relevance beyond traditional epilepsy syndromes.^86^ This study therefore lays the groundwork for future research to assess whether actively mitigating sleep fragmentation can suppress epileptiform discharges, lower seizure risk, and minimise disruptions to memory processing and cognitive function associated with nocturnal interictal activity.

Our study was designed to include a heterogeneous patient population to identify consistent effects of sleep stability on epileptic activity across diverse clinical presentations, enhancing the generalisability of our findings. Consequently, the cohort included a broad range of structural abnormalities, as well as MRI-negative cases, and histopathological confirmation was not available for all participants. The resulting heterogeneity and small numbers within individual pathological categories precluded meaningful statistical subgroup analyses to determine whether the observed regional effects varied according to the underlying lesion or histopathological substrate. As such, drawing definitive conclusions about specific epilepsy subtypes or pathological substrates was beyond the scope of this work. Nevertheless, our findings provide a valuable foundation for future multicentre investigations in larger cohorts, with sufficient representation of specific pathological substrates, to determine whether the regional modulation of arousal-induced IEDs varies according to the underlying pathology and to further delineate subtype-specific mechanisms. The SOZ was defined for each patient using clinical electrographic criteria based on the earliest detected ictal changes and therefore may not correspond to the true epileptogenic zone. The SOZ should thus be considered a functional electrographic construct that includes regions involved in seizure initiation. Another limitation is that this study was conducted in patients with drug-resistant epilepsy, the only population in whom intracranial electrodes can be implanted inside the human brain for clinical purposes. As such, the findings may not generalise to individuals with controlled epilepsy. The influence of implantation density on our findings was reduced by restricting our main analyses to active channels with IEDs and adopting a channel-level GLMM approach. However, implantation strategy remains clinically driven, and the resulting set of active channels may not sample the underlying epileptic network uniformly or reflect the degree to which sampled tissue is irritative or lesional. Future studies combining channel-level activity measures with tissue-level pathological correlates will help to further resolve this distinction. Finally, while our study design allowed us to probe the effects of arousals on IEDs more directly than prior observational studies, we acknowledge that IEDs can also precipitate arousals, indicating a bidirectional relationship, and that underlying brain states may influence this interaction.^19^ Future studies should combine behavioural testing with simultaneous sleep and EEG recordings to clarify the functional consequences of arousal-driven IED increases on memory and cognition, paving the way for interventions that mitigate the neurological consequences of sleep disruption in epilepsy.

In conclusion, our study provides novel evidence that transient sleep instability, specifically evoked arousals induced during NREM sleep, can acutely amplify interictal activity in drug-resistant epilepsy. By experimentally manipulating sleep stability with precise temporal control, we demonstrate that arousal-driven interictal spiking is anatomically and functionally selective, occurring in the neocortex and with a larger effect during N2 sleep. Further, we show that while arousals increase interictal spike burden, they do not influence spatial propagation. This suggests that arousal-related spiking reflects heightened local cortical excitability without sufficient recruitment of large-scale epileptic networks, offering important mechanistic insights into the distinct processes underlying the generation and spread of epileptic phenomena. Together, these findings reinforce the potential of sleep-based paradigms as a powerful tool to probe network dynamics in epilepsy as well as other pathological brain states. Our work also highlights sleep as a modifiable therapeutic target for neuromodulatory approaches, with the potential to reduce interictal and ictal burden and improve cognitive outcomes in people with epilepsy.

## Methods

### Patient selection

Adult patients with drug-resistant focal epilepsy undergoing SEEG as part of their clinical presurgical evaluation were recruited. A total of 20 patients were enrolled between May 2021 and January 2025 across two sites: the Montreal Neurological Institute and Hospital (MNI) and the Duke Comprehensive Epilepsy Centre (DCEC). Three patients were excluded from the analysis: two due to technical issues with the auditory stimulation protocol, and one due to abnormal sleep patterns that made sleep scoring unreliable. The final cohort included 17 patients (8 females, 14 right-handed, mean age of 37.2 ± 8.1; **Supplementary Table 1**), with 10 patients from the MNI and 7 from the DCEC. The study was approved by the MNI Review Ethics Board (2021-7516) and the Duke University Health System Institutional Review Board (Pro00113635), and informed written consent was obtained from each patient.

### SEEG-PSG recordings

All participants underwent combined SEEG-PSG recordings as described in previous studies.^11,12,19,30^ SEEG depth electrodes were implanted intracranially to localise the epileptogenic zone, with electrode placement determined by clinical teams at the MNI and DCEC. SEEG signals were sampled at 2048 Hz, referenced to an electrode positioned over the parietal lobe contralateral to the suspected epileptogenic zone, using the Neuroworkbench system (Nihon Kohden, Tokyo, Japan) at the MNI or the Natus Quantum LTM amplifier at DCEC (Natus Medical Inc., Middleton, WI, USA). For patients at the MNI, SEEG signals were high-pass filtered at 0.08 Hz and low-pass filtered at 600 Hz. At the DCEC, signals were high-pass filtered at 0.01 Hz and low-pass filtered at 1000 Hz. PSG recordings and the auditory stimulation protocol were performed early during the presurgical evaluation, at least 48 hours after implantation of SEEG electrodes and while patients remained either on their full antiseizure medication regimen or had undergone only a limited, clinically indicated medication reduction as part of their routine care. The PSG recordings included scalp EEG from standard 10-20 system sites (F3,F4,C3,C4,O1,O2,M1,M2) obtained with subdermal thin wire electrodes placed during implantation,^87^ along with surface electrooculography (EOG) and electromyography (EMG) of the chin and tibialis anterior muscles to facilitate sleep scoring. No seizures were observed during the auditory stimulation sessions analysed.

### Auditory stimulation protocol

The objective of the auditory stimulation protocol was to induce arousals, following guidelines from the American Sleep Disorders Association.^88^ Auditory stimuli consisted of pure tones with a frequency of 1 kHz, lasting 5 seconds, and ranging in volume from 40 to 100 dB. Stimuli were manually delivered via high-fidelity tubal insert earphones (ER3C, Etyomic, Illinois, USA) during the first half of the night defined as 4 hours starting from the participant’s habitual bedtime, enabling precise sound delivery while minimising external noise interference and contamination in EEG recordings. Continuous online PSG monitoring was performed by the experimenter trained in sleep scoring (SH, AH) to assess sleep stage in real time and to guide stimulus delivery, including confirming whether an evoked arousal was successfully induced. This real-time assessment was based on visual inspection of ongoing EEG, EOG and EMG activity, according to standard sleep scoring criteria from the American Academy of Sleep Medicine (AASM).^13^ Stimulation began after 5 minutes of stable NREM sleep, marked by the appearance of a sleep spindle or K-complex,^13^ and sound volume was gradually increased until an arousal occurred. The stimulation protocol was applied during both NREM and REM sleep to maximise the occurrence of sleep arousals during real-time sleep scoring; however, analyses focused on NREM sleep due to its well-established role in facilitating epileptic activity, and because many patients did not reach REM sleep due to the stimulation-induced sleep fragmentation. If stimulation caused an awakening, the protocol was stopped and restarted at a lower volume (-10 dB) after 2 minutes of stable sleep, defined by the reappearance of a sleep spindle, K-complex, or rapid eye movements in the PSG recordings. In cases where stimulation induced an arousal, the next stimulus was delivered at the same volume after 2 minutes of stable sleep. If no arousal occurred, the next stimulus was administered after 10 seconds at a higher volume (+10 dB). This protocol has been shown to increase the arousal index by up to 20 events/hour without affecting total sleep time.^31,89^

### Sleep scoring and arousal detection

Sleep stages and arousals were scored manually in accordance with the standard criteria of the American Academy of Sleep Medicine (AASM).^13^ Scoring was based on PSG recordings, including scalp EEG, EOG, and EMG, and was performed by trained personnel blinded to the SEEG data (SH, LP-D). Sleep was scored in 30-second epochs; each epoch was classified into one of the standard sleep stages (N1, N2, N3, REM, or wake) based on characteristic EEG, EOG, and EMG patterns. An arousal was defined as an abrupt shift of EEG frequency including alpha, theta and/or frequencies greater than 16 Hz (not including sleep spindles) that lasted 3-15 s, with ≥10 s of stable sleep preceding the change.^13^ Arousals were classified according to the sleep stage in which they occurred. Arousals occurring within 6.5 s of auditory stimulus onset were considered evoked. This window accounted for the 5-second duration of stimulus presentation and an additional 1.5 s to allow for the delay between stimulation application and arousal manifestation.^90^ Arousals falling outside this window were classified as spontaneous arousals and were subsequently analysed to determine whether findings are specific to auditory evoked arousals or generalise to spontaneous arousals. All sleep stage and arousal scoring were verified by a board-certified neurologist with subspecialty training in sleep medicine (BF, LP-D).

### Epoch selection

To assess the impact of arousals on IEDs, we analysed four distinct 3-second epochs in the SEEG that were time-locked to each auditory stimulation event in all patients (**Fig. 1c**). These time windows were chosen to capture transient changes in interictal activity occurring immediately before and after auditory stimulation and arousal onset. A 3-second duration provided a sufficient temporal resolution to detect rapid changes in IEDs while minimising overlap between successive events and preserving temporal specificity. This epoch duration was chosen specifically to capture the local, immediate sleep microstate surrounding each auditory stimulus and arousal event, rather than longer-timescale sleep dynamics, and also aligns with previous work examining peri-arousal neural dynamics.^12,19^ For auditory stimuli that successfully induced arousals, we extracted a pre-stimulus baseline (the 3-second epoch immediately preceding stimulus onset) and an evoked arousal window (the 3-second epoch starting at arousal onset). To control for the effects of auditory stimulation in the absence of arousals, we identified two additional 3-second control epochs surrounding auditory stimuli that did not induce arousals: a control pre-stimulus baseline, defined as the 3 seconds immediately before these non-arousing stimuli, and a control 3-s epoch beginning 1.08 s after stimulus onset, defined as the 3-second epoch beginning 1.08 s after the non-arousing stimulus onset, corresponding to the median stimulus-to-arousal onset latency across all evoked arousals (IQR = 0.46-2.10 s, range = 0.02-6.33 s). This design allowed us to isolate the specific effects of arousals on interictal activity, while controlling for non-specific effects of auditory input. These carefully matched epochs provided the basis for quantifying the effects of transient sleep disruptions on both IED occurrence and spatial propagation, enabling a precise characterisation of arousal-related epileptic dynamics. For analysis of spontaneous arousals, a 3-s epoch beginning at arousal onset and a corresponding 3-s pre-arousal baseline epoch immediately preceding arousal onset were selected for comparison with control epochs.

### Analysis of interictal epileptiform discharges

IEDs were automatically detected within the selected epochs on bipolar SEEG channels using a previously validated detector with default parameters,^40^ which has been independently verified by our group and utilised in several studies.^11,30,91–93^. To account for inter-individual differences in electrode implantation density, analyses were restricted to active channels, defined as those with an IED rate exceeding 4.4 events/min, calculated across all analysed epochs and corresponding to the false positive rate of the IED detector used.^40^ Channel-level IED counts in active SEEG channels were subsequently modelled using a generalised linear mixed-effects model (GLMM) (see ‘Statistical analyses’). To examine the spatial propagation of IEDs, we quantified IED propagation by calculating the number of propagation channels, defined as the number of SEEG channels recruited during each independent IED.^11,30^ IEDs were considered independent if they occurred at least 120 ms apart from a preceding IED on any channel.^11,30^ The first detected IED in a given epoch was considered the source IED, and any additional spikes across all SEEG channels within 120 ms of the source IED were considered propagating IEDs. IED count per channel and number of propagating channels were computed separately for all four pre-defined epochs including the arousal conditions (pre-stimulus baseline and arousal window for auditory stimuli that induced arousals) and the control conditions (pre-stimulus baseline and latency-matched control window for non-arousing stimuli). HFOs were also detected automatically on bipolar SEEG channels within the selected epochs, using a previously validated detector,^94^ to identify HFO events (>80 Hz). Channel-level HFO counts in active SEEG channels were subsequently modelled using a generalised linear mixed-effects model (GLMM) (see ‘Statistical analyses’).

### Analysis of modulating factors

To examine the factors modulating arousal-related changes in interictal activity, we assessed the influence of anatomical brain region, involvement of the seizure-onset zone (SOZ), and sleep stage. This enabled analysis of network-level differences in arousal-driven interictal activity. Since we did not observe a significant effect of arousals on IED propagation, subsequent analysis of modulating factors was performed only on IED counts. Electrode localisation was performed following established co-registration procedures used in our previous work.^95,96^ Post-implantation CT/MRI scans displaying electrode positions were co-registered to each patient’s pre-implantation MRI images, which were in turn non-linearly aligned to the ICBM152 non-linear symmetric brain model.^97^ This combined transformation enabled electrode positions to be accurately determined within a common stereotaxic space. Electrode channels were subsequently assigned to anatomical regions based on the MICCAI atlas using a 5-mm radius sphere centred on each contact coordinate.^95^ Channels were categorised as mesiotemporal (including amygdala, hippocampus, and entorhinal cortex) or neocortical, to reflect known differences in functional connectivity with the thalamus, a key regulator of sleep.^42^ Neocortical regions receive extensive thalamocortical projections whereas mesiotemporal regions are more sparsely connected to the thalamus.^42,57^ This anatomical categorisation was consistent with previous SEEG studies characterising sleep-epilepsy interactions.^11,19^ IED counts were computed separately for source IEDs originating in neocortical and mesiotemporal regions.

The influence of the seizure-onset zone (SOZ) on arousal-related interictal dynamics were examined by categorising IEDs according to whether they originated within or outside the SOZ. The SOZ was identified for each patient by a board-certified neurologist with subspecialty training and certification in clinical neurophysiology and epilepsy (BF), based on clinical SEEG recordings. Specifically, seizure onset was defined as the earliest sustained rhythmic EEG change that was visually distinguishable from background activity, in line with established criteria.^30,43^ The SOZ was defined as the set of SEEG contacts exhibiting this activity at seizure onset. Where multiple seizures were recorded, SOZ classification was based on consistent involvement of contacts across seizures. SOZ designation was determined through visual inspection of SEEG traces in conjunction with clinical reports, reflecting standard presurgical evaluation procedures.^98^ Each SEEG channel was labelled as either ‘SOZ’ or ‘non-SOZ’, enabling separate quantification of IED counts across the four 3-s epochs associated with evoked arousal and control conditions, based on involvement of SOZ networks. Patients with less than three SEEG channels in any channel subset (mesiotemporal, neocortical, SOZ, or non-SOZ) were excluded from the corresponding analyses.

To assess whether arousal-related changes in interictal activity were dependent on sleep stage, we separated out events based on the NREM sleep stage in which they were delivered. Specifically, auditory stimuli were grouped based on whether they occurred during epochs marked as stage N2 or stage N3 sleep, using standard sleep scoring criteria.^13^ Arousals overlapping stage transitions were excluded from this stage-specific analysis to ensure stable sleep state classification. IED counts across all SEEG channels were then calculated separately for each sleep stage across evoked arousal and control conditions. Patients who did not have evoked arousals or corresponding control epochs during either N2 or N3 sleep were excluded from this analysis. This approach allowed us to evaluate sleep stage-dependent susceptibility of interictal networks to evoked arousals, providing valuable insights into whether arousal-driven interictal spiking is modulated by the underlying sleep architecture.

To assess the influence of slow-wave activity on arousal-related IED modulation, slow waves were automatically detected on bipolar SEEG channels using an algorithm previously implemented and validated in-house.^45^ Stimulus responses were classified as either arousals with slow-wave activity or isolated slow-wave responses according to whether a slow wave was detected on any channel during the 3-s arousal epoch or latency-matched control epoch, respectively.

### Statistical analyses

All IED metrics, namely IED count and number of propagation channels, were compared across the four pre-defined 3-second epochs of interest: the pre-stimulus baseline preceding auditory stimuli that induced evoked arousals, the evoked arousal window, the pre-stimulus baseline for non-arousing stimuli (control), and the latency-matched control window for non-arousing stimuli. These 3-s epochs allowed us to capture local sleep microstates immediately before the auditory stimulus and during the arousal, consistent with a prior SEEG study assessing IEDs during sleep.^19^ To confirm that IED counts during the 3-s pre-stimulus epochs were representative of the patient’s overall baseline, we compared normalised IED counts between pre-stimulus epochs and randomly selected arousal-free epochs of equivalent duration sampled across the recording night, using a GLMM with a Tweedie distribution and a log link function, with epoch type included as a fixed effect and patient ID as a random effect.

IED count was modelled using a GLMM with a negative binomial distribution and a log link function, appropriate for modelling overdispersed count data such as IED counts. Condition (evoked arousal vs control) was included as the fixed effect of primary interest, with pre-stimulus baseline IED count and stimulus intensity included as fixed-effect covariates, and random intercepts for SEEG contact, nested within patient ID, to account for between-contact and between-patient variability in baseline IED rates. Mean propagation channel numbers across IEDs were modelled using a Gamma GLMM with a log link function, appropriate for positively skewed, continuous response variables. Condition (evoked arousal vs control) was included as the fixed effect of primary interest, with pre-stimulus number of propagation channels and stimulus intensity included as fixed-effect covariates, and a random intercept for patient ID. For the main evoked arousal analysis, four IED count models were fitted: one across all active channels, and three further models each including an interaction term between condition and a subgroup variable representing each modulating factor of interest: anatomical brain region (mesiotemporal vs neocortical), SOZ involvement (inside vs outside SOZ), and sleep stage (N2 vs N3 sleep) to formally test whether the effect of evoked arousal differed according to each modulating factor. As the propagation channel model for the main evoked arousal analysis did not yield a significant effect of condition, no further analysis of propagation-related modulating factors was performed.

To evaluate whether these findings generalised to spontaneous arousals, equivalent negative binomial and Gamma GLMM architectures were implemented to assess IED counts and IED propagation, respectively, during spontaneous arousals, adjusted for baseline IED count. Additionally, HFO counts during evoked arousals were modelled using a negative binomial GLMM, adjusted for baseline IED count and stimulus intensity, to determine whether the effect of arousals on IEDs extended to other markers of epileptogenicity. For both spontaneous arousals and HFOs, identical interaction terms (anatomical region, SOZ status, and sleep stage) were incorporated to examine potential modulating factors.

To assess whether pre-existing differences in baseline brain state could account for the observed increase in IED counts during evoked arousals, pre-stimulus IED count was modelled using an additional GLMM with a negative binomial distribution and log link function, with condition (evoked arousal vs control) included as the fixed effect of primary interest, stimulus intensity included as a fixed-effect covariate, and random intercepts for SEEG contact, nested within patient ID. We also assessed whether the acute effects of evoked arousals on IEDs extended to a larger temporal scale by comparing global, patient-level IED rates during NREM sleep between the first half of the night, during which auditory stimulation was delivered to induce sleep fragmentation, and the second half of the night, during which the stimulation protocol was withheld to allow recovery sleep. This first-half/second-half design was adopted a priori to allow comparison of periods of high and low sleep fragmentation while keeping antiseizure medication levels and seizure state closely matched within the same patient and night. Global IED rates during the fragmentation and recovery periods were compared using a two-sided Wilcoxon signed-rank test. This pattern was further assessed using randomly sampled, arousal-free 3-s baseline epochs across both halves of the night, modelled using a GLMM with a Tweedie distribution and a log link function, with epoch type (’stim on’ vs ‘stim off’) included as a fixed effect and patient ID as a random effect.

To directly test whether the pre-stimulus background state conditioned the electrophysiological response to stimulation, two further sets of models were fitted. Pre-stimulus spectral power ratio was defined as δ/(θ+α+β) ([0.5-4 Hz]/[4.5-30 Hz]), derived from the scalp EEG (Fz-Cz, or F3-C3 in two patients where Fz-Cz was unavailable).^19^ This ratio was calculated over the 3-s epoch immediately preceding the auditory stimulus, then log10-transformed and z-scored. Power values were derived within standard EEG frequency bands (δ: 0.5–4 Hz; θ: 4.5–8 Hz; α: 8–12 Hz; β: 12–30 Hz).^19^ First, the probability of an arousal response, relative to an isolated slow-wave response, to stimulation was modelled using a binomial GLMM with a logit link function, with pre-stimulus spectral power ratio, stimulus intensity, and pre-stimulus baseline IED count included as fixed-effect covariates, and random intercepts for SEEG contact, nested within patient ID, to account for between-contact and between-patient variability. This model tested whether the pre-stimulus delta activity, as measured by the spectral power ratio, predicted the probability of an arousal response to stimulation. Secondly, to test whether pre-stimulus delta state modulated the relationship between stimulus response type and epileptiform activity, IED count was modelled using a negative binomial GLMM with a log link function, including response type (isolated slow-wave response vs arousal response), pre-stimulus spectral power ratio, and their interaction as fixed effects, alongside pre-stimulus baseline IED count and stimulus intensity as covariates, with random intercepts for SEEG contact nested within patient ID. As the ‘response type × pre-stimulus spectral power ratio’ interaction reached significance, post-hoc simple slopes analysis was performed to estimate the slope of pre-stimulus spectral power ratio on IED count separately for each response type, to characterise the direction and significance of this relationship and thereby clarify the ‘permissive’ versus ‘protective’ role of pre-stimulus delta activity. To address the possibility that pre-stimulus delta state represented a direct confounding pathway between response type and IED count, a change-in-estimate sensitivity analysis was performed: the effect of response type on IED count was estimated in an unadjusted negative binomial GLMM (excluding pre-stimulus spectral power ratio) and compared with an adjusted model additionally including this covariate.

To address the possibility that stimulus intensity itself was driving IED counts, IED count during control epochs was modelled using a negative binomial GLMM with a log link function, with stimulus intensity as the fixed effect of interest, adjusting for pre-stimulus baseline IED count, and including random intercepts for SEEG contact nested within patient ID. This model tested whether stimulus intensity was associated with IED count in the absence of a detectable arousal response. We then tested whether the difference in IED count between evoked arousals and control epochs depended on stimulus intensity, by modelling IED count across all evoked arousal and control epochs using a negative binomial GLMM with a log link function, including event type (evoked arousal vs control), stimulus intensity, and their interaction as fixed effects, adjusting for pre-stimulus baseline IED count, and including random intercepts for SEEG contact nested within patient ID. To test whether stimulus intensity was associated with changes in thalamic spectral content during control epochs, the thalamic spectral power ratio – δ/(θ+α+β) ([0.5-4 Hz]/[4.5-30 Hz]) – was computed from bipolar-referenced thalamic contacts in the four patients who had SEEG electrodes implanted within the thalamus.^19^ The change in thalamic spectral power ratio, calculated as the difference between the control and pre-stimulus epochs, was modelled using a Gaussian-family GLMM with stimulus intensity as the fixed effect of interest, and random intercepts for SEEG contact nested within patient ID.

To control for multiple comparisons, models were grouped into pre-specified families defined by outcome variable (IED count, HFO count, and propagation channel number) and analysis context (evoked arousal, spontaneous arousal, and slow-wave activity). We applied a two-tiered correction approach. First, the *p*-values for the fixed effect (or interaction term) of primary interest were Bonferroni-corrected across the number of models within each family. Specifically, the main evoked arousal IED count analysis constituted one family of four models (*p* × 4); similarly, the spontaneous arousal IED count and evoked arousal HFO count analyses each constituted separate families of four models (*p* × 4); the slow-wave analysis family consisted of two models (*p* × 2); and the propagation channel models for evoked and spontaneous arousals were treated as separate single-model families. Secondly, for any model in which the interaction term reached significance, *p*-values for the associated pairwise post-hoc contrasts were further Bonferroni-corrected within model (*p* × 2): this applied both to comparisons of arousal vs control within each subgroup level in the main evoked arousal, spontaneous arousal and HFO analyses, and to the simple slopes comparison of isolated slow wave vs arousal in the slow-wave analysis. Effect sizes for fixed effects are reported as incidence rate ratios (IRR) and model-predicted means are presented with 95% confidence intervals.

To characterise the temporal evolution of interictal spiking around arousal and stimulus onset, IED counts for each patient were normalised to that patient’s maximum value, aggregated into 500 ms bins, and averaged across patients with group data presented as mean ± standard error. IED count data from all SEEG channels exhibiting IEDs were included. For each post-onset bin, group-level normalised IED counts were compared against the mean pre-onset baseline values across patients using one-tailed Mann– Whitney tests. Statistically significant bins (*p* < 0.05) are indicated by asterisks in the corresponding plots. To examine the relationship between arousal duration and IED rates, we used a generalised linear mixed-effects model (GLMM) with a Gamma distribution and a log link function, appropriate for modelling positively skewed, continuous response variables such as IED rates. The independent variable was arousal duration (in seconds), and the dependent variable was either the normalised IED count (total number of IEDs during the full arousal per channel) or IED rate (number of IEDs per second), across the full arousal duration. Arousals with undefined normalised IED values due to a baseline value of zero were excluded from the GLMM analysis. To account for inter-individual variability and repeated measures within patients, patient ID was included as a random intercept.

We assessed whether thalamic spectral activity was associated with arousal-related changes in IED count by conducting a subset analysis in the four patients who had SEEG electrodes implanted within the thalamus. A total of 56 evoked arousals occurring during N2 sleep were included across these patients. For each arousal, the thalamic spectral power ratio – δ/(θ+α+β) ([0.5-4 Hz]/[4.5-30 Hz]) – was computed from bipolar-referenced thalamic contacts during the 3-s arousal epoch.^19^ This ratio, reflecting the relative dominance of slow versus fast thalamic activity, was normalised to the corresponding 3-s pre-stimulus baseline epoch. IED counts were extracted from all SEEG channels during the same 3-s arousal epoch, and normalised both to the number of SEEG channels and to the pre-stimulus baseline. We then applied a GLMM with a Gamma distribution and log link to test whether thalamic spectral power ratio predicted normalised IED count. The model included thalamic spectral power ratio as a fixed effect and a random intercept for patient ID to account for repeated measures within individuals. Model estimates were expressed as β ± standard error (SE), and statistical significance was assessed using Wald tests (*P* < 0.05). All GLMM results are presented as fixed-effect coefficient estimates (β-estimate ± standard error), alongside *p*-values, with 95% confidence intervals indicated in the corresponding plots. For comparisons of global IED rate and arousal rate at the patient level between stimulation-induced sleep fragmentation and recovery sleep, effect sizes were calculated using Cliff’s delta (Cliff’s *d*) to quantify the magnitude of observed differences. All statistical analyses were conducted using a significance threshold of *α* = 0.05.

## Data availability

This study is based on clinical data collected under institutional ethics committee approval, with informed consent obtained from all participants. Data will be made available upon reasonable request, subject to compliance with the conditions of the ethical approval governing its collection and use. Source data for the main Results and Supplementary Results are provided with this paper.

## Code availability

For scoring of sleep and arousals, we used Stellate Harmonie (Stellate, Montreal, QC, Canada). Custom Python scripts were developed to compute IED counts and propagation channel numbers as well as HFO counts, extract sleep and evoked arousal annotations, and stratify outputs by channel subset and sleep stage (available on our GitHub repository at https://github.com/Lab-Frauscher/Auditory_Stimulation). These scripts utilise functions from the EPYCOM repository (version 0.3, 2025; https://doi.org/10.5281/zenodo.18193447),^99^ which includes previously developed code for IED detection and processing, based on the methods described by Janca et al. (2015),^40^ and for HFO detection, based on methods described by von Ellenrieder et al. (2016).^94^ Code for slow-wave detection is available on our Gitlab repository (https://gitlab.com/anphy_duke/lab-tools/slow-wave-detector), based on methods described by Frauscher et al (2015).^45^ All statistical analyses were performed using MATLAB 2023b (MathWorks, Santa Clara, CA, USA), R (version 4.4.1, R Core Team, 2024; https://www.r-project.org/), and Jamovi (version 2.6, The Jamovi Project, 2025; https://www.jamovi.org/).

## Acknowledgements

We express our gratitude to the staff and technicians in the EEG Department of the Montreal Neurological Institute and Hospital (in particular, Lorraine Allard, Nicole Drouot, and Chantal Lessard) and the Duke Comprehensive Epilepsy Centre (in particular, Mary Byrd, Emily Kale, Crystal Keller, and Jasmine Patterson). We also thank Tanguy Hedrich for his technical support in setting up the auditory stimulation protocol. This work was supported by a Postdoctoral Fellowship from the Fonds de Recherche du Québec–Santé (FRQS) (2021-2023), as well as a Professional Development Award and start-up funding from the Faculty of Health and Medicine, Lancaster University, all held by SH. BF was supported by a project grant from the Canadian Institutes of Health Research (PJT-175056) and start-up funding from Duke University.

## Contributions

Study design and conceptualisation: B.F., L.P-D., and S.H. Data acquisition: A.H., E.M., and S.H. Data analysis: B.M., K.J., T.A., J.T., D.C., P.K., L.P-D., and S.H. Interpretation: B.F., L.P-D., and S.H. Writing – original draft: S.H. Writing – Review & Editing: B.F., A.H., B.M., K.J., T.A., E.M., J.T., D.S., J.H., D.C., P.K., L.P-D., and S.H. Resources: B.F., D.S., and J.H. Funding acquisition: B.F. and S.H.

## References

1 Sheybani, L., Frauscher, B., Bernard, C. & Walker, M. C. Mechanistic insights into the interaction between epilepsy and sleep. Nat Rev Neurol 21, 177–192 (2025). 10.1038/s41582-025-01064-z

2 Latreille, V., St Louis, E. K. & Pavlova, M. Co-morbid sleep disorders and epilepsy: A narrative review and case examples. Epilepsy Res 145, 185–197 (2018). 10.1016/j.eplepsyres.2018.07.005

3 Nobili, L. et al. Sleep and epilepsy: A snapshot of knowledge and future research lines. J Sleep Res 31, e13622 (2022). 10.1111/jsr.13622

4 Latreille, V. et al. Nocturnal seizures are associated with more severe hypoxemia and increased risk of postictal generalized EEG suppression. Epilepsia 58, e127–e131 (2017). 10.1111/epi.13841

5 Gibbs, E. & Gibbs, F. Diagnostic and localizing value of electroencephalographic studies in sleep| CiNii Research. Res Publ Assoc Res Nerv Ment Dis 26, 366–376 (1947).

6 Janz, D. The grand mal epilepsies and the sleeping-waking cycle. Epilepsia 3, 69–109 (1962). 10.1111/j.1528-1157.1962.tb05235.x

7 Halász, P. Sleep and epilepsy. Handb Clin Neurol 107, 305–322 (2012). 10.1016/b978-0-444-52898-8.00019-7

8 Halász, P., Terzano, M., Parrino, L. & Bódizs, R. The nature of arousal in sleep. J Sleep Res 13, 1–23 (2004). 10.1111/j.1365-2869.2004.00388.x

9 Herman, S. T., Walczak, T. S. & Bazil, C. W. Distribution of partial seizures during the sleep--wake cycle: differences by seizure onset site. Neurology 56, 1453–1459 (2001). 10.1212/wnl.56.11.1453

10 Ng, M. & Pavlova, M. Why are seizures rare in rapid eye movement sleep? Review of the frequency of seizures in different sleep stages. Epilepsy Res Treat 2013, 932790 (2013). 10.1155/2013/932790

11 Ho, A. et al. Rapid eye movement sleep affects interictal epileptic activity differently in mesiotemporal and neocortical areas. Epilepsia 64, 3036–3048 (2023). 10.1111/epi.17763

12 Wang, Y. L. et al. Intracerebral Dynamics of Sleep Arousals: A Combined Scalp-Intracranial EEG Study. J Neurosci 44 (2024). 10.1523/jneurosci.0617-23.2024

13 Troester, M. M., Quan, S. F. & Berry, R. B. The AASM Manual for the Scoring of Sleep and Associated Events: Rules, Terminology and Technical Specicifications. Version 3. (American Academy of Sleep Medicine, 2023).

14 Manni, R., Zambrelli, E., Bellazzi, R. & Terzaghi, M. The relationship between focal seizures and sleep: an analysis of the cyclic alternating pattern. Epilepsy Res 67, 73–80 (2005). 10.1016/j.eplepsyres.2005.08.008

15 Giorgi, F. S. et al. Cyclic alternating pattern and interictal epileptiform discharges during morning sleep after sleep deprivation in temporal lobe epilepsy. Epilepsy Behav 73, 131–136 (2017). 10.1016/j.yebeh.2017.05.005

16 Parrino, L., Halasz, P., Tassinari, C. A. & Terzano, M. G. CAP, epilepsy and motor events during sleep: the unifying role of arousal. Sleep Med Rev 10, 267–285 (2006). 10.1016/j.smrv.2005.12.004

17 Malow, A., Bowes, R. J. & Ross, D. Relationship of temporal lobe seizures to sleep and arousal: a combined scalp-intracranial electrode study. Sleep 23, 231–234 (2000).

18 Terzaghi, M. et al. Sleep-related minor motor events in nocturnal frontal lobe epilepsy. Epilepsia 48, 335–341 (2007). 10.1111/j.1528-1167.2006.00929.x

19 Peter-Derex, L. et al. Sleep Disruption in Epilepsy: Ictal and Interictal Epileptic Activity Matter. Ann Neurol 88, 907–920 (2020). 10.1002/ana.25884

20 Burlando, G. et al. Unstable slow oscillations couple with epileptogenic fast-rhythm bistability in sleep-related epilepsy: A stereoelectroencephalographic study. Epilepsia (2026). 10.1002/epi.70188

21 Zubler, F., Rubino, A., Lo Russo, G., Schindler, K. & Nobili, L. Correlating Interictal Spikes with Sigma and Delta Dynamics during Non-Rapid-Eye-Movement-Sleep. Front Neurol 8, 288 (2017). 10.3389/fneur.2017.00288

22 Frauscher, B. & Gotman, J. Sleep, oscillations, interictal discharges, and seizures in human focal epilepsy. Neurobiol Dis 127, 545–553 (2019). 10.1016/j.nbd.2019.04.007

23 Hannan, S., Ho, A. & Frauscher, B. Clinical Utility of Sleep Recordings During Presurgical Epilepsy Evaluation With Stereo-Electroencephalography: A Systematic Review. J Clin Neurophysiol 41, 430–443 (2024). 10.1097/wnp.0000000000001057

24 Jayakar, P. et al. Diagnostic utility of invasive EEG for epilepsy surgery: Indications, modalities, and techniques. Epilepsia 57, 1735–1747 (2016). 10.1111/epi.13515

25 Kang, X. et al. Quantitative spatio-temporal characterization of epileptic spikes using high density EEG: Differences between NREM sleep and REM sleep. Sci Rep 10, 1673 (2020). 10.1038/s41598-020-58612-4

26 McLeod, G. A., Ghassemi, A. & Ng, M. C. Can REM Sleep Localize the Epileptogenic Zone? A Systematic Review and Analysis. Front Neurol 11, 584 (2020). 10.3389/fneur.2020.00584

27 Grigg-Damberger, M. & Foldvary-Schaefer, N. Bidirectional relationships of sleep and epilepsy in adults with epilepsy. Epilepsy Behav 116, 107735 (2021). 10.1016/j.yebeh.2020.107735

28 Lambert, I. et al. Brain regions and epileptogenicity influence epileptic interictal spike production and propagation during NREM sleep in comparison with wakefulness. Epilepsia 59, 235–243 (2018). 10.1111/epi.13958

29 Parvizi, J. & Kastner, S. Promises and limitations of human intracranial electroencephalography. Nat Neurosci 21, 474–483 (2018). 10.1038/s41593-018-0108-2

30 Hannan, S. et al. The Differing Effects of Sleep on Ictal and Interictal Network Dynamics in Drug-Resistant Epilepsy. Ann Neurol (2023). 10.1002/ana.26796

31 Martin, S. E., Engleman, H. M., Deary, I. J. & Douglas, N. J. The effect of sleep fragmentation on daytime function. Am J Respir Crit Care Med 153, 1328–1332 (1996). 10.1164/ajrccm.153.4.8616562

32 Lambert, I. et al. Hippocampal Interictal Spikes during Sleep Impact Long-Term Memory Consolidation. Ann Neurol 87, 976–987 (2020). 10.1002/ana.25744

33 Camarillo-Rodriguez, L. et al. Temporal lobe interictal spikes disrupt encoding and retrieval of verbal memory: A subregion analysis. Epilepsia 63, 2325–2337 (2022). 10.1111/epi.17334

34 Holmes, G. L. Interictal Spikes as an EEG Biomarker of Cognitive Impairment. J Clin Neurophysiol 39, 101–112 (2022). 10.1097/wnp.0000000000000728

35 Xu, Y. et al. Impact of interictal epileptiform discharges on brain network in self-limited epilepsy with centrotemporal spikes: A magnetoencephalography study. Brain Behav 13, e3038 (2023). 10.1002/brb3.3038

36 Cheng, D., Yan, X., Xu, K., Zhou, X. & Chen, Q. The effect of interictal epileptiform discharges on cognitive and academic performance in children with idiopathic epilepsy. BMC Neurol 20, 233 (2020). 10.1186/s12883-020-01807-z

37 Rasch, B. & Born, J. About sleep’s role in memory. Physiol Rev 93, 681–766 (2013). 10.1152/physrev.00032.2012

38 Keeble, L., Monaghan, P., Robertson, E. M. & Hannan, S. Slow-wave sleep as a key player in offline memory processing: insights from human EEG studies. Front Behav Neurosci 19, 1620544 (2025). 10.3389/fnbeh.2025.1620544

39 Adamantidis, A. R., Gutierrez Herrera, C. & Gent, T. C. Oscillating circuitries in the sleeping brain. Nat Rev Neurosci 20, 746–762 (2019). 10.1038/s41583-019-0223-4

40 Janca, R. et al. Detection of interictal epileptiform discharges using signal envelope distribution modelling: application to epileptic and non-epileptic intracranial recordings. Brain Topogr 28, 172–183 (2015). 10.1007/s10548-014-0379-1

41 Chiang, S. et al. State-dependent effects of responsive neurostimulation depend on seizure localization. Brain 148, 521–532 (2025). 10.1093/brain/awae240

42 Gent, T. C., Bassetti, C. & Adamantidis, A. R. Sleep-wake control and the thalamus. Curr Opin Neurobiol 52, 188–197 (2018). 10.1016/j.conb.2018.08.002

43 Spanedda, F., Cendes, F. & Gotman, J. Relations between EEG seizure morphology, interhemispheric spread, and mesial temporal atrophy in bitemporal epilepsy. Epilepsia 38, 1300–1314 (1997). 10.1111/j.1528-1157.1997.tb00068.x

44 Bernard, C., Frauscher, B., Gelinas, J. & Timofeev, I. Sleep, oscillations, and epilepsy. Epilepsia 64 Suppl 3, S3–s12 (2023). 10.1111/epi.17664

45 Frauscher, B. et al. Facilitation of epileptic activity during sleep is mediated by high amplitude slow waves. Brain 138, 1629–1641 (2015). 10.1093/brain/awv073

46 Prince, D. A. & Connors, B. W. Mechanisms of interictal epileptogenesis. Adv Neurol 44, 275–299 (1986).

47 Avoli, M. Mechanisms of epileptiform synchronization in cortical neuronal networks. Curr Med Chem 21, 653–662 (2014). 10.2174/0929867320666131119151136

48 Lambert, I. et al. Cortico-cortical and thalamo-cortical connectivity during non-REM and REM sleep: Insights from intracranial recordings in humans. Clin Neurophysiol 143, 84–94 (2022). 10.1016/j.clinph.2022.08.026

49 Timofeev, I., Bazhenov, M., Seigneur, J. & Sejnowski, T. in Jasper’s Basic Mechanisms of the Epilepsies (eds J. L. Noebels et al.) (National Center for Biotechnology Information (US) Copyright © 2012, Michael A Rogawski, Antonio V Delgado-Escueta, Jeffrey L Noebels, Massimo Avoli and Richard W Olsen., 2012).

50 Peter-Derex, L., Magnin, M. & Bastuji, H. Heterogeneity of arousals in human sleep: A stereo-electroencephalographic study. Neuroimage 123, 229–244 (2015). 10.1016/j.neuroimage.2015.07.057

51 Halász, P., Kelemen, A. & Szűcs, A. The role of NREM sleep micro-arousals in absence epilepsy and in nocturnal frontal lobe epilepsy. Epilepsy Res 107, 9–19 (2013). 10.1016/j.eplepsyres.2013.06.021

52 Tomlinson, S. B., Wong, J. N., Conrad, E. C., Kennedy, B. C. & Marsh, E. D. Reproducibility of interictal spike propagation in children with refractory epilepsy. Epilepsia 60, 898–910 (2019). 10.1111/epi.14720

53 Tomlinson, S. B. et al. Spatiotemporal Mapping of Interictal Spike Propagation: A Novel Methodology Applied to Pediatric Intracranial EEG Recordings. Front Neurol 7, 229 (2016). 10.3389/fneur.2016.00229

54 Maharathi, B. et al. Interictal spike connectivity in human epileptic neocortex. Clin Neurophysiol 130, 270–279 (2019). 10.1016/j.clinph.2018.11.025

55 McCormick, D. A. & Bal, T. Sleep and arousal: thalamocortical mechanisms. Annu Rev Neurosci 20, 185–215 (1997). 10.1146/annurev.neuro.20.1.185

56 Jones, E. G. The thalamic matrix and thalamocortical synchrony. Trends Neurosci 24, 595–601 (2001). 10.1016/s0166-2236(00)01922-6

57 Ketz, N. A., Jensen, O. & O’Reilly, R. C. Thalamic pathways underlying prefrontal cortex-medial temporal lobe oscillatory interactions. Trends Neurosci 38, 3–12 (2015). 10.1016/j.tins.2014.09.007

58 Colrain, I. M. & Campbell, K. B. The use of evoked potentials in sleep research. Sleep Med Rev 11, 277–293 (2007). 10.1016/j.smrv.2007.05.001

59 Kamali, A. et al. The Cortico-Limbo-Thalamo-Cortical Circuits: An Update to the Original Papez Circuit of the Human Limbic System. Brain Topogr 36, 371–389 (2023). 10.1007/s10548-023-00955-y

60 Aggleton, J. P. et al. Hippocampal-anterior thalamic pathways for memory: uncovering a network of direct and indirect actions. Eur J Neurosci 31, 2292–2307 (2010). 10.1111/j.1460-9568.2010.07251.x

61 Van der Werf, Y. D., Witter, M. P. & Groenewegen, H. J. The intralaminar and midline nuclei of the thalamus. Anatomical and functional evidence for participation in processes of arousal and awareness. Brain Res Brain Res Rev 39, 107–140 (2002). 10.1016/s0165-0173(02)00181-9

62 Fouad, A. et al. Interictal epileptiform discharges show distinct spatiotemporal and morphological patterns across wake and sleep. Brain Commun 4, fcac183 (2022). 10.1093/braincomms/fcac183

63 Lundstrom, B. N. & Richner, T. J. Neural adaptation and fractional dynamics as a window to underlying neural excitability. PLoS Comput Biol 19, e1010527 (2023). 10.1371/journal.pcbi.1010527

64 Yang, Y., Zhang, F., Gao, X., Feng, L. & Xu, K. Progressive alterations in electrophysiological and epileptic network properties during the development of temporal lobe epilepsy in rats. Epilepsy Behav 141, 109120 (2023). 10.1016/j.yebeh.2023.109120

65 Hatlestad-Hall, C. et al. Source-level EEG and graph theory reveal widespread functional network alterations in focal epilepsy. Clin Neurophysiol 132, 1663–1676 (2021). 10.1016/j.clinph.2021.04.008

66 Parrino, L., Ferri, R., Bruni, O. & Terzano, M. G. Cyclic alternating pattern (CAP): the marker of sleep instability. Sleep Med Rev 16, 27–45 (2012). 10.1016/j.smrv.2011.02.003

67 Terzano, M. G. & Parrino, L. Origin and Significance of the Cyclic Alternating Pattern (CAP). REVIEW ARTICLE. Sleep Med Rev 4, 101–123 (2000). 10.1053/smrv.1999.0083

68 Mak-McCully, R. A. et al. Coordination of cortical and thalamic activity during non-REM sleep in humans. Nat Commun 8, 15499 (2017). 10.1038/ncomms15499

69 Bello, S. T. et al. Visually or auditorily induced seizures involve the activation of nonhippocampal brain areas and hippocampal removal does not alleviate seizures in a mouse model of temporal lobe epilepsy. Epilepsia 65, 218–237 (2024). 10.1111/epi.17816

70 Quimby, A. E. et al. Signal processing and stimulation potential within the ascending auditory pathway: a review. Front Neurosci 17, 1277627 (2023). 10.3389/fnins.2023.1277627

71 Baxendale, S. & Heaney, D. Memory complaints in the epilepsy clinic. Pract Neurol (2020). 10.1136/practneurol-2020-002523

72 Tinson, D. et al. Memory complaints in epilepsy: An examination of the role of mood and illness perceptions. Epilepsy Behav 80, 221–228 (2018). 10.1016/j.yebeh.2017.11.028

73 Mazarati, A. Epilepsy and forgetfulness: one impairment, multiple mechanisms. Epilepsy Curr 8, 25–26 (2008). 10.1111/j.1535-7511.2007.00224.x

74 Ackermann, S. & Rasch, B. Differential effects of non-REM and REM sleep on memory consolidation? Curr Neurol Neurosci Rep 14, 430 (2014). 10.1007/s11910-013-0430-8

75 Brodt, S., Inostroza, M., Niethard, N. & Born, J. Sleep-A brain-state serving systems memory consolidation. Neuron 111, 1050–1075 (2023). 10.1016/j.neuron.2023.03.005

76 Lambert, I. et al. Accelerated long-term forgetting in focal epilepsy: Do interictal spikes during sleep matter? Epilepsia 62, 563–569 (2021). 10.1111/epi.16823

77 Karoly, P. J. et al. Interictal spikes and epileptic seizures: their relationship and underlying rhythmicity. Brain 139, 1066–1078 (2016). 10.1093/brain/aww019

78 de Curtis, M. & Avanzini, G. Interictal spikes in focal epileptogenesis. Prog Neurobiol 63, 541–567 (2001). 10.1016/s0301-0082(00)00026-5

79 Avoli, M. et al. Network and pharmacological mechanisms leading to epileptiform synchronization in the limbic system in vitro. Prog Neurobiol 68, 167–207 (2002). 10.1016/s0301-0082(02)00077-1

80 Rehim, E. D., Vendrame, M. & Devinsky, O. Obstructive sleep apnea in people with epilepsy: Modifying risk. Epilepsia (2026). 10.1002/epi.70375

81 Liguori, C., Toledo, M. & Kothare, S. Effects of anti-seizure medications on sleep architecture and daytime sleepiness in patients with epilepsy: A literature review. Sleep Med Rev 60, 101559 (2021). 10.1016/j.smrv.2021.101559

82 Ren, G. et al. Impact of antiseizure medication taper on electroencephalographic dynamics in focal epilepsy: A stereoelectroencephalographic study. Epilepsia 67, 2585–2600 (2026). 10.1002/epi.70149

83 Wei, Y., Krishnan, G. P., Marshall, L., Martinetz, T. & Bazhenov, M. Stimulation Augments Spike Sequence Replay and Memory Consolidation during Slow-Wave Sleep. J Neurosci 40, 811–824 (2020). 10.1523/jneurosci.1427-19.2019

84 Ngo, H. V., Martinetz, T., Born, J. & Mölle, M. Auditory closed-loop stimulation of the sleep slow oscillation enhances memory. Neuron 78, 545–553 (2013). 10.1016/j.neuron.2013.03.006

85 Zeller, C. J., Züst, M. A., Wunderlin, M., Nissen, C. & Klöppel, S. The promise of portable remote auditory stimulation tools to enhance slow-wave sleep and prevent cognitive decline. J Sleep Res 32, e13818 (2023). 10.1111/jsr.13818

86 Lemus, H. N. & Sarkis, R. A. Interictal epileptiform discharges in Alzheimer’s disease: prevalence, relevance, and controversies. Front Neurol 14, 1261136 (2023). 10.3389/fneur.2023.1261136

87 Ives, J. R. New chronic EEG electrode for critical/intensive care unit monitoring. J Clin Neurophysiol 22, 119–123 (2005). 10.1097/01.wnp.0000152659.30753.47

88 EEG arousals: scoring rules and examples: a preliminary report from the Sleep Disorders Atlas Task Force of the American Sleep Disorders Association. Sleep 15, 173–184 (1992).

89 Martin, S. E., Brander, P. E., Deary, I. J. & Douglas, N. J. The effect of clustered versus regular sleep fragmentation on daytime function. J Sleep Res 8, 305–311 (1999). 10.1046/j.1365-2869.1999.00169.x

90 Bastuji, H., Perchet, C., Legrain, V., Montes, C. & Garcia-Larrea, L. Laser evoked responses to painful stimulation persist during sleep and predict subsequent arousals. Pain 137, 589–599 (2008). 10.1016/j.pain.2007.10.027

91 Klimes, P. et al. NREM sleep is the state of vigilance that best identifies the epileptogenic zone in the interictal electroencephalogram. Epilepsia 60, 2404–2415 (2019). 10.1111/epi.16377

92 Azeem, A. et al. Interictal spike networks predict surgical outcome in patients with drug-resistant focal epilepsy. Ann Clin Transl Neurol 8, 1212–1223 (2021). 10.1002/acn3.51337

93 Klimes, P., Peter-Derex, L., Hall, J., Dubeau, F. & Frauscher, B. Spatio-temporal spike dynamics predict surgical outcome in adult focal epilepsy. Clin Neurophysiol 134, 88–99 (2022). 10.1016/j.clinph.2021.10.023

94 von Ellenrieder, N., Frauscher, B., Dubeau, F. & Gotman, J. Interaction with slow waves during sleep improves discrimination of physiologic and pathologic high-frequency oscillations (80-500 Hz). Epilepsia 57, 869–878 (2016). 10.1111/epi.13380

95 Frauscher, B. et al. Atlas of the normal intracranial electroencephalogram: neurophysiological awake activity in different cortical areas. Brain 141, 1130–1144 (2018). 10.1093/brain/awy035

96 Avigdor, T. et al. The Awakening Brain is Characterized by a Widespread and Spatiotemporally Heterogeneous Increase in High Frequencies. Adv Sci (Weinh) 12, e2409608 (2025). 10.1002/advs.202409608

97 Fonov, V. et al. Unbiased average age-appropriate atlases for pediatric studies. Neuroimage 54, 313–327 (2011). 10.1016/j.neuroimage.2010.07.033

98 Frauscher, B. et al. Learn how to interpret and use intracranial EEG findings. Epileptic Disord 26, 1–59 (2024). 10.1002/epd2.20190

99 Cimbalnik, J. et al. EPYCOM - ElectroPhYsiology COmputational Module (Version 0.3) (2025). Zenodo. 10.5281/zenodo.18193447

