## Supplemental Material for "Sleep fragmentation drives local, network-specific epileptic activity in the human brain"

### Supplementary Table 1. Patient Demographics

| Patient ID | Age | Sex | Handedness | Number of implanted electrodes | Type of epilepsy | Epileptic focus | MRI lesion | Anti-seizure medication during SEEG investigation |
|---|---|---|---|---|---|---|---|---|
| 1 | 41 | M | R | 14 (L) | TLE | L mesiotemporal | smaller L hippocampus and unusual gyral formation of L fusiform gyrus | carbamazepine, levetiracetam |
| 2 | 27 | F | L | 14 (R) | ETLE | R latero-occipital | no abnormality | lacosamide, levetiracetam, |
| 3 | 44 | F | R | 16 (R) | ETLE | R posterio-lateral parietal | no abnormality | carbamazepine, lacosamide, levetiracetam, phenytoin |
| 4 | 46 | M | R | 12 (2 R, 10 L) | ETLE | L parieto-occipital | L parieto-occipital lesion | carbamazepine, clobazam, lacosamide, lamotrigine, lorazepam |
| 5 | 40 | F | L | 15 (L) | TLE | L mesiotemporal | L temporal atrophy (mesial and lateral) extending to insula and perisylvian area | clobazam, lacosamide, lamotrigine, topiramate |
| 6 | 33 | M | R | 13 (R) | TLE | R posterior temporal/posterior insula | R temporal neocortex atrophy atrophy/agenesis of the R piriform | brivaracetam, clonazepam, eslicarbazepine, lacosamide, pregabalin |
| 7 | 29 | M | R | 13 (R) | ETLE | R posterior | no abnormality | carbamazepine, levetiracetam, phenobarbital |
| 8 | 37 | F | R | 10 (L) | TLE | L mesiobasal temporal | mild L mesiotemporal sclerosis | carbamazepine, lamotrigine |
| 9 | 36 | F | R | 9 (R) | TLE+ | R temporo-insular | T2 flair hypersensitivity in the residual mesial/lateral temporal lobe (gliotic changes) | escitalopram, topiramate |
| 10 | 41 | M | R | 11 (R) | ETLE | R posterior quadrant | PNH, R atrium lateral ventricle, remote R anterior temporal lobe resection, stable encephalomalacic changes along surgical borders | clobazam, lacosamide, oxcarbazepine |
| 11 | 38 | M | R | 18 (L) | TLE | L temporal opercular region/superior temporal gyrus | no abnormality | brivaracetam, clonazepam, lacosamide, midazolam |
| 12 | 28 | F | R | 18 (L) | ETLE | L posterior cortex | L middle cranial fossa arachnoid cyst | clobazam, oxcarbazepine |
| 13 | 22 | F | R | 16 (L) | TLE | Lfusiform gyrus in close vicinity to remote resection | remote L posterior basal temporal ganglioglioma resection | cenobamate, lamotrigine, zonisamide |
| 14 | 41 | F | R | 19 (16 L, 3 R) | TLE | L temporal pole and amygdala | L hippocampal sclerosis | cenobamate, clobazam, levetiracetam, lacosamide |
| 15 | 53 | M | R | 20 (12 L, 8 R) | ETLE | L insula in close vicinity of cavernoma | cavernoma in L insula | clobazam, lacosamide, perampanel |
| 16 | 37 | M | L | 16 (L) | TLE+ | L mesial orbitofrontal, L mesiotemporal | questionable FCD in L frontobasal region. | brivaracetam, cenobamate, lamotrigine |
| 17 | 45 | M | R | 17 (15 L, 2 R) | TLE | L mesiotemporal, L temporal pole | PNH within L occipital/frontal lobe. Possible L hippocampal atrophy | carbamazepine, diazepam, levetiracetam |

[a]In any patient who had two or more distinct epileptic foci, seizures that arose from the same focus were selected from the two states for analysis.

Abbreviations: F = female; M = male; R = right; L = left; TLE = temporal lobe epilepsy; TLE+ = temporal lobe epilepsy with additional involvement of other brain regions; ETLE = extratemporal lobe epilepsy; FCD = focal cortical dysplasia; PNH = periventricular nodular heterotopia.

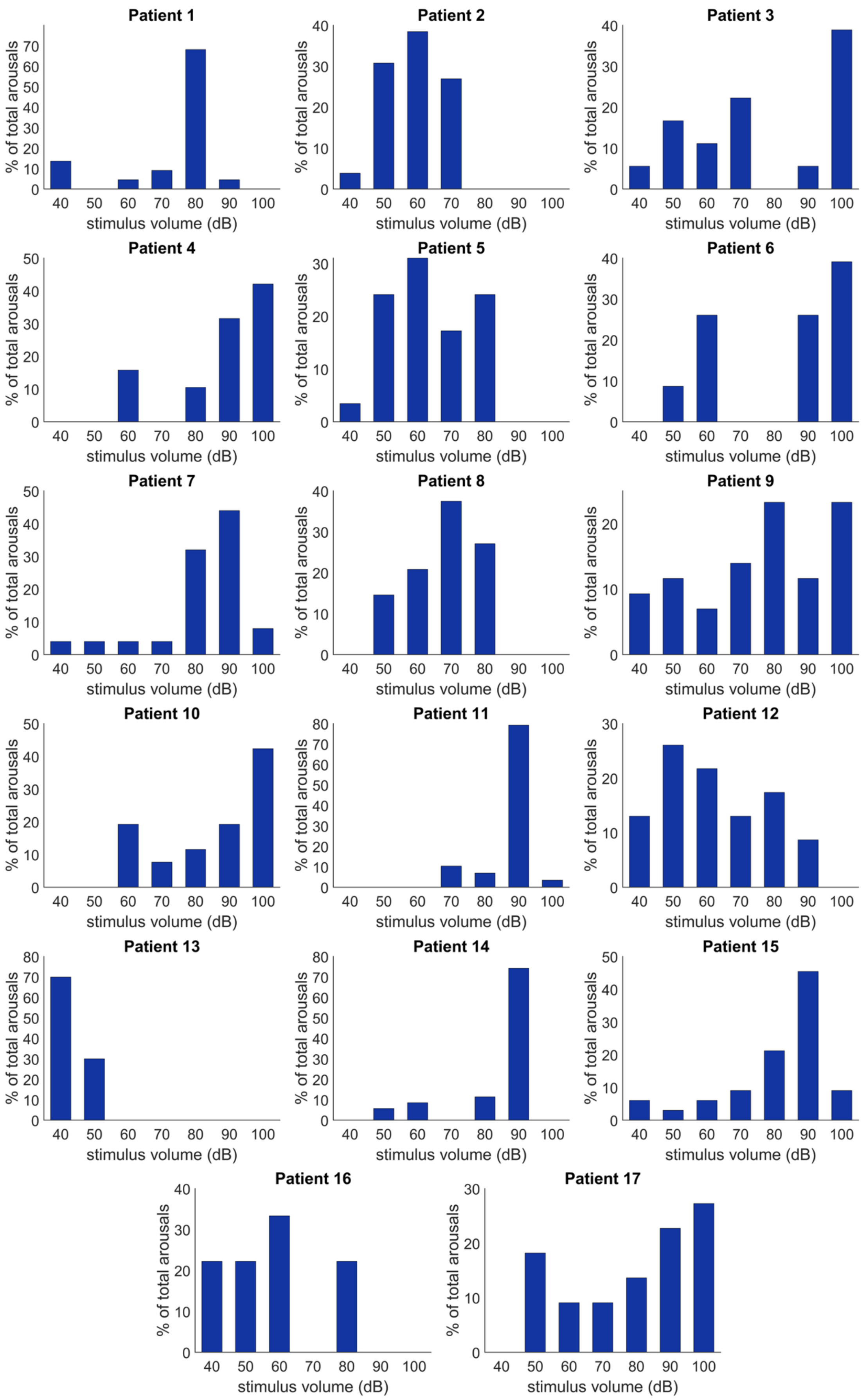


**Supplementary Figure 1. Patient-level distribution of evoked arousals across auditory stimulus intensities.** The percentage of total evoked arousals induced by each stimulus intensity (dB) is shown for each patient in our cohort (*n* = 17). Each value represents the proportion of arousals elicited at a given intensity relative to that patient's total number of evoked arousals. Patients demonstrated variable sensitivity to auditory stimulation, with some showing robust responses at lower intensities (e.g., Patients 2, 5, and 13 at 40-60 dB), while others required higher intensities (≥80 dB) to reliably evoke arousals (e.g., Patients 1, 4, 7, 11, and 14). This heterogeneity reflects individual differences in auditory arousal thresholds.

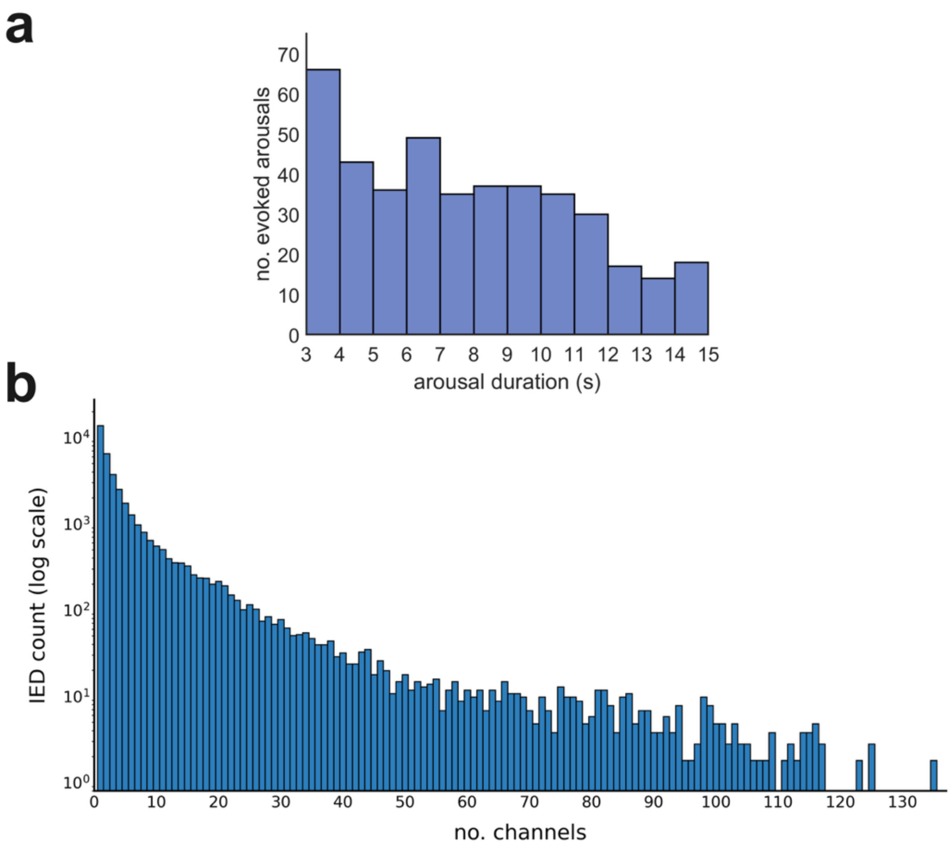


**Supplementary Figure 2. Distribution of arousal durations and SEEG channel involvement in interictal epileptiform discharges (IEDs). (a)** Histogram of the durations (in seconds) of arousals evoked by auditory stimulation (bin width: 1 s). **(b)** Log-scale histogram of the number of SEEG channels involved in each IED (bin width: 1 channel). Histograms show total numbers of arousals and IEDs used in analysis, pooled across all patients.

**Supplementary Table 2. Global interictal epileptiform discharge (IED) counts and rates per patient across the night.**

| Patient ID | No. auditory stimulations | Global IED count | Global IED rate (events/min) | IED rate/channel (events/min) |
|---|---|---|---|---|
| 1 | 74 | 22,035 | 91.8 | 0.70 |
| 2 | 67 | 70,100 | 140.2 | 0.98 |
| 3 | 99 | 46,492 | 85.8 | 0.84 |
| 4 | 68 | 34,977 | 89.8 | 0.59 |
| 5 | 74 | 26,358 | 52.1 | 0.47 |
| 6 | 96 | 15,546 | 34.5 | 0.22 |
| 7 | 99 | 23,437 | 79.4 | 0.53 |
| 8 | 58 | 26,597 | 69.8 | 0.74 |
| 9 | 82 | 19,679 | 51.4 | 0.56 |
| 10 | 105 | 50,502 | 121.4 | 1.03 |
| 11 | 92 | 29,665 | 72.2 | 0.35 |
| 12 | 85 | 67,849 | 148.8 | 0.74 |
| 13 | 68 | 37,078 | 99.0 | 0.50 |
| 14 | 126 | 51,509 | 98.8 | 0.49 |
| 15 | 146 | 38,081 | 177.5 | 0.94 |
| 16 | 84 | 32,339 | 82.8 | 0.42 |
| 17 | 110 | 68,334 | 197.5 | 1.11 |

For each patient: total number of auditory stimulations delivered during the recording session, total IED count captured across all sleep during the night, global IED rate, and IED rate per channel. IEDs were identified across all SEEG channels exhibiting IEDs during the night, with events considered independent if they occurred at least 120 ms apart from a preceding IED on any channel.

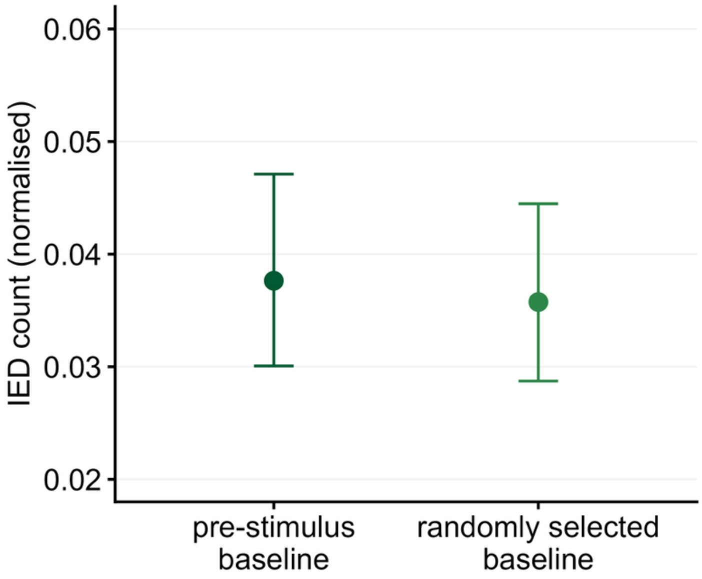


**Supplementary Figure 3. Pre-stimulus baseline interictal epileptiform discharge (IED) counts are comparable to IED counts sampled from baseline arousal-free epochs across the night.** Model-predicted mean of the IED count, normalised by the number of active SEEG channels, for the 3-s pre-stimulus baseline epochs used in the main analysis, compared with randomly selected arousal-free epochs of equivalent duration sampled across the recording night. IED counts from pre-stimulus baseline epochs did not differ significantly to those from randomly selected baseline epochs (generalised linear mixed-effects model (GLMM), β-estimate ± standard error (SE): -0.052 ± 0.027, $P$ = 0.053), indicating that the pre-stimulus epochs are representative of the underlying baseline IED rate across the night. Error bars represent 95% confidence intervals.

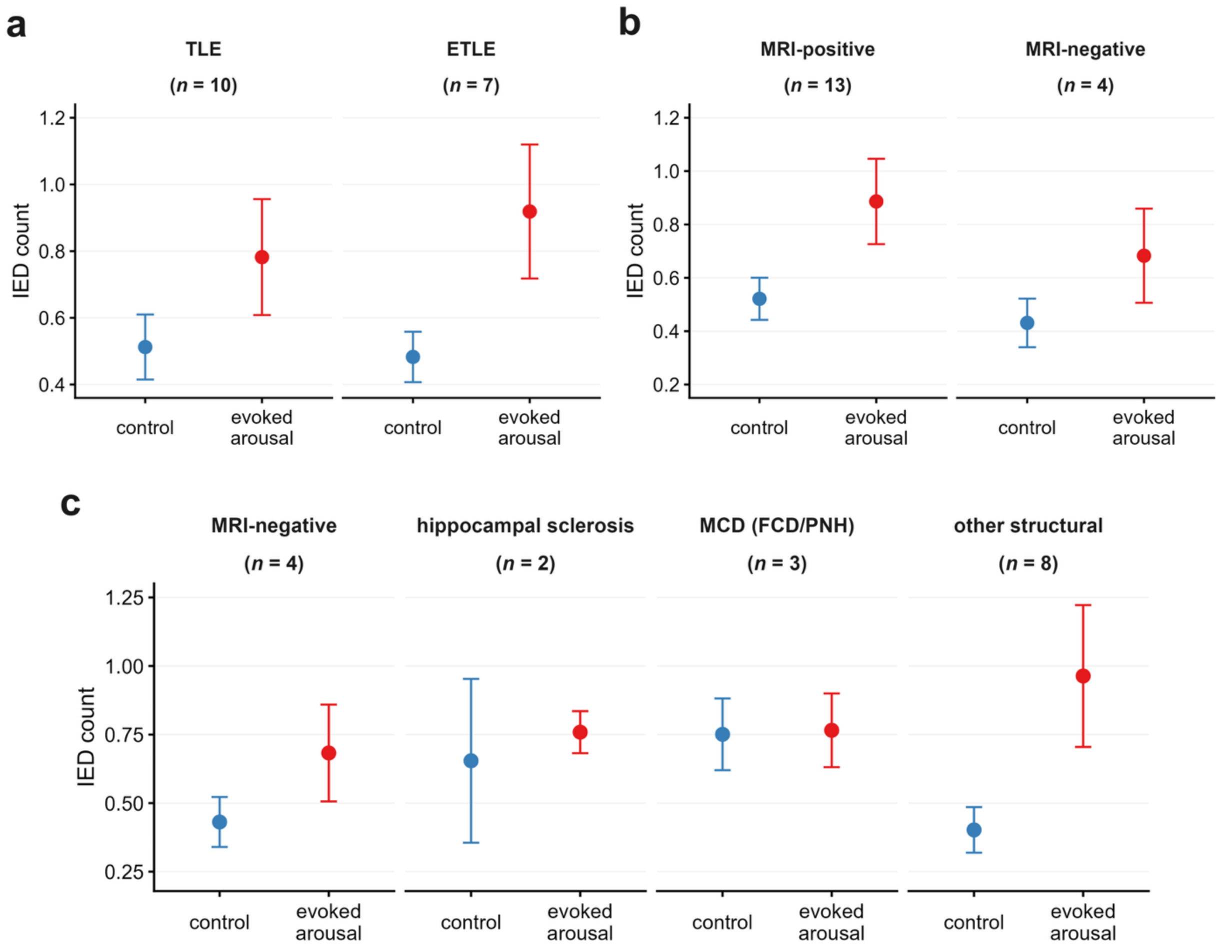


**Supplementary Figure 4. Exploratory subgroup comparisons of arousal-related interictal epileptiform discharge (IED) counts by epilepsy type and MRI status.** Descriptive plots showing mean IED count (± SE) during control (blue) and evoked arousal (red) epochs, stratified by **(a)** epilepsy type (temporal lobe epilepsy [TLE] vs extratemporal lobe epilepsy [ETLE]), **(b)** MRI status (MRI-positive vs MRI-negative), and **(c)** MRI lesion category (MRI-negative, hippocampal sclerosis, malformations of cortical development (MCD, including focal cortical dysplasia [FCD] and periventricular nodular heterotopia [PNH]), and other structural pathology (temporal/hippocampal atrophy, gliosis, arachnoid cyst, cavernoma, and ganglioglioma resection)). Circles and error bars depict group mean ± standard error (SE), calculated across patient averages (mean IED count per active SEEG channel per event epoch (control vs evoked arousal)). Patient sample sizes per category are indicated in parentheses (*n*).

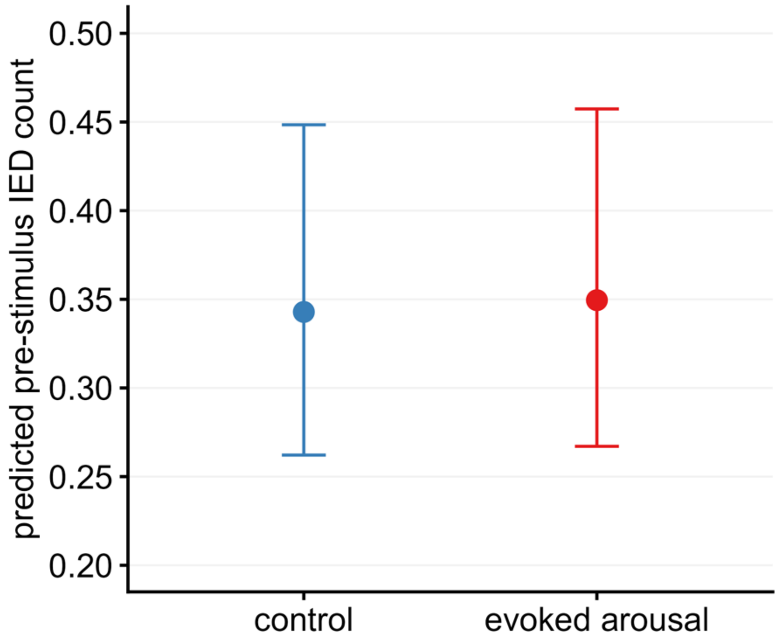


**Supplementary Figure 5. Pre-stimulus interictal epileptiform discharge (IED) counts do not differ between control and evoked arousal conditions.** Model-predicted mean IED count for the 3-s pre-stimulus epochs preceding control trials, compared with those preceding evoked arousals, adjusting for stimulus intensity. IED counts from pre-stimulus epochs preceding evoked arousals did not differ significantly from those preceding controls (generalised linear mixed-effects model (GLMM), β-estimate ± standard error (SE): 0.019 ± 0.014, incidence rate ratio (IRR) = 1.02, *P* = 0.182), indicating that the increase in IED count observed during evoked arousals is not attributable to a pre-existing difference in baseline IED rate between conditions. Error bars represent 95% confidence intervals.

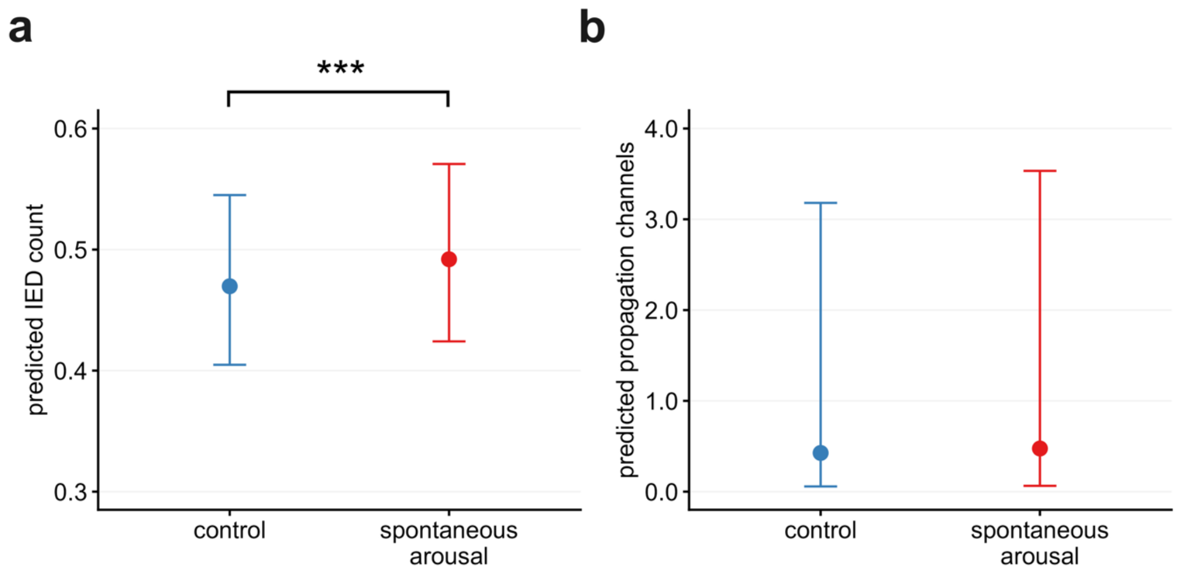


**Supplementary Figure 6. Spontaneous arousals increase interictal epileptiform discharge (IED) counts but not propagation. (a)** Model-predicted mean IED counts per channel-epoch under control and spontaneous arousal conditions, adjusted for baseline IED count. Predicted IED count was significantly higher during spontaneous arousals than controls, corresponding to a 5% increase (GLMM, β-estimate ± standard error (SE): 0.046 ± 0.010 SE, incidence rate ratio: 1.05, $p < 0.0001$). **(b)** Model-predicted IED propagation channel numbers per epoch under control and spontaneous arousal conditions, adjusted for baseline propagation channel number. The number of propagation channels did not differ significantly between spontaneous arousals and controls (GLMM, β-estimate ± SE: 0.109 ± 0.317 SE, IRR: 1.12, $P = 0.733$). Error bars represent 95% confidence intervals. ***$p < 0.001$.

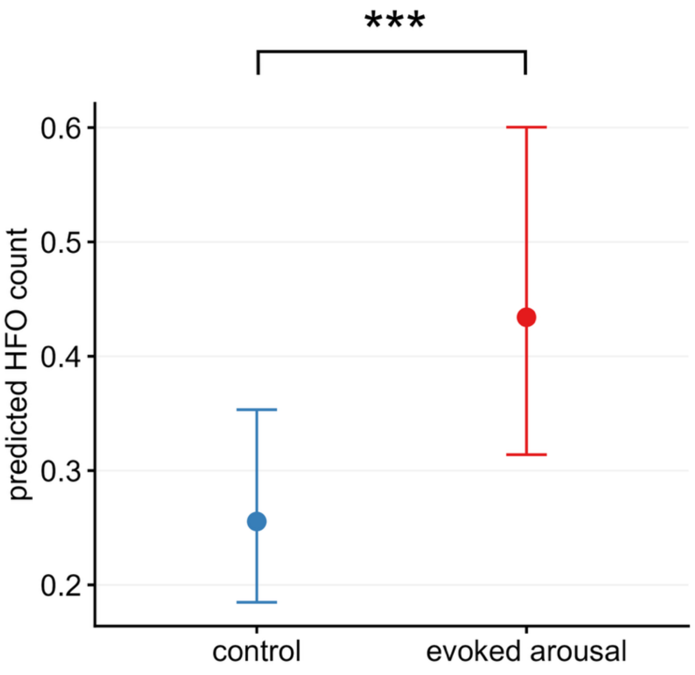


**Supplementary Figure 7. Evoked arousals increase high-frequency oscillations (HFOs).** Model-predicted mean HFO counts per channel-epoch under control and evoked arousal conditions, adjusted for pre-stimulus HFO count and stimulus intensity. Predicted HFO count was significantly higher during evoked arousals than controls, corresponding to a 70% increase (GLMM, β-estimate ± standard error (SE): 0.530 ± 0.014 SE, IRR: 1.70, $p < 0.0001$). Error bars represent 95% confidence intervals. ***$p < 0.001$.

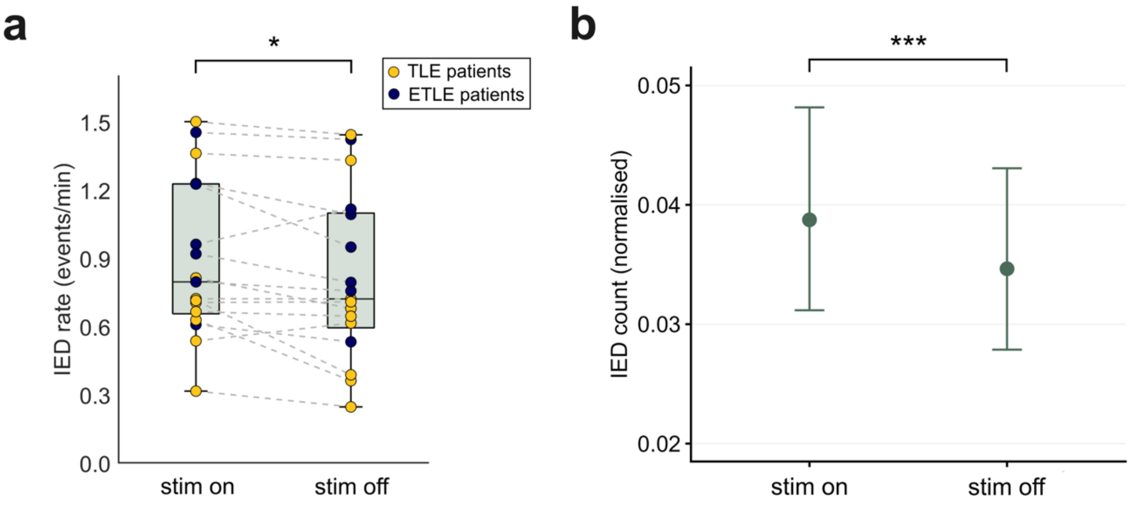


**Supplementary Figure 8. Global interictal epileptiform discharge (IED) rates are elevated during stimulation-induced sleep fragmentation. (a)** Global patient-level IED rates during NREM sleep, normalised by active SEEG channels, during the first half of the sleep night, when the auditory stimulation protocol was delivered to induce sleep fragmentation ('stim on'), and the second half of the night when auditory stimulation was withheld, allowing recovery sleep ('stim off'). IED rates were significantly higher during stimulation-induced sleep fragmentation than periods without auditory stimulation (Wilcoxon signed rank test, $p = 0.017$, Cliff's $d = 0.15$). Data are presented as individual patient values. Patients with temporal lobe epilepsy (TLE; yellow circles) and extratemporal lobe epilepsy (ETLE; navy blue circles) are distinguished for visual comparison. No consistent differences in global IED rates between TLE and ETLE subgroups were observed. **(b)** Model-predicted mean normalised IED count from randomly sampled 3-s baseline epochs during 'stim on' and 'stim off' periods across the night. IED count was significantly higher during stimulation-induced sleep fragmentation than during periods without stimulation (generalised linear mixed-effects model (GLMM), β-estimate ± standard error (SE): -0.112 ± 0.003, rate ratio = 0.89, $p < 0.0001$). Error bars represent 95% confidence intervals. *$p < 0.05$, ***$p < 0.001$.

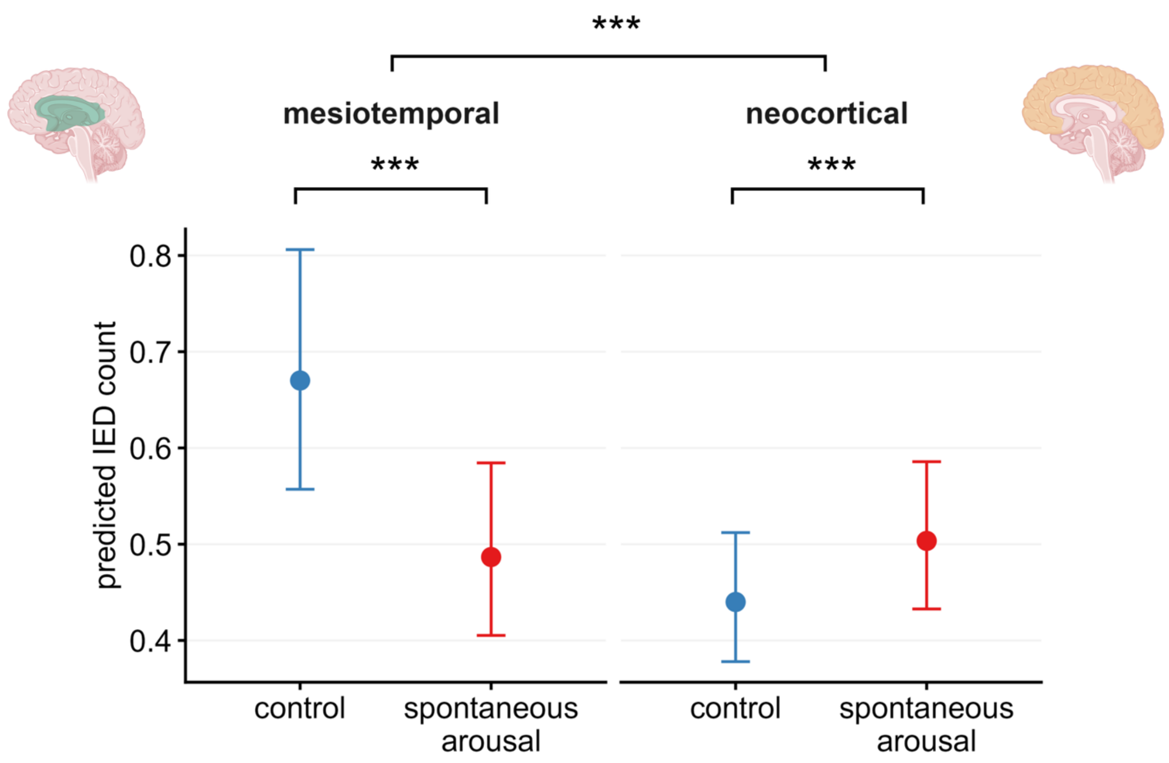


**Supplementary Figure 9. Spontaneous arousals are associated with divergent effects on interictal epileptiform discharges (IEDs) in mesiotemporal and neocortical regions.** Model-predicted mean IED counts per channel-epoch by anatomical region (mesiotemporal vs neocortical) under control and spontaneous arousal conditions, adjusted for baseline IED count. The interaction between condition and anatomical region was significant (generalised linear mixed-effects model (GLMM), β-estimate ± standard error (SE): 0.455 ± 0.030, incidence rate ratio (IRR) = 1.58, $p < 0.0001$), indicating that the effect of evoked arousals on IED count differed between mesiotemporal and neocortical regions. In mesiotemporal regions ($n$ = 14 patients), predicted IED count showed a significant 27% decrease during spontaneous arousals relative to controls (β = -0.320 ± 0.028, IRR = 0.73, $p < 0.0001$). In neocortical regions ($n$ = 17 patients), predicted IED count was significantly higher during spontaneous arousals than controls (β = 0.135 ± 0.011, IRR = 1.14, $p < 0.0001$), corresponding to a 14% increase. Error bars represent 95% confidence intervals. *** $p < 0.001$.

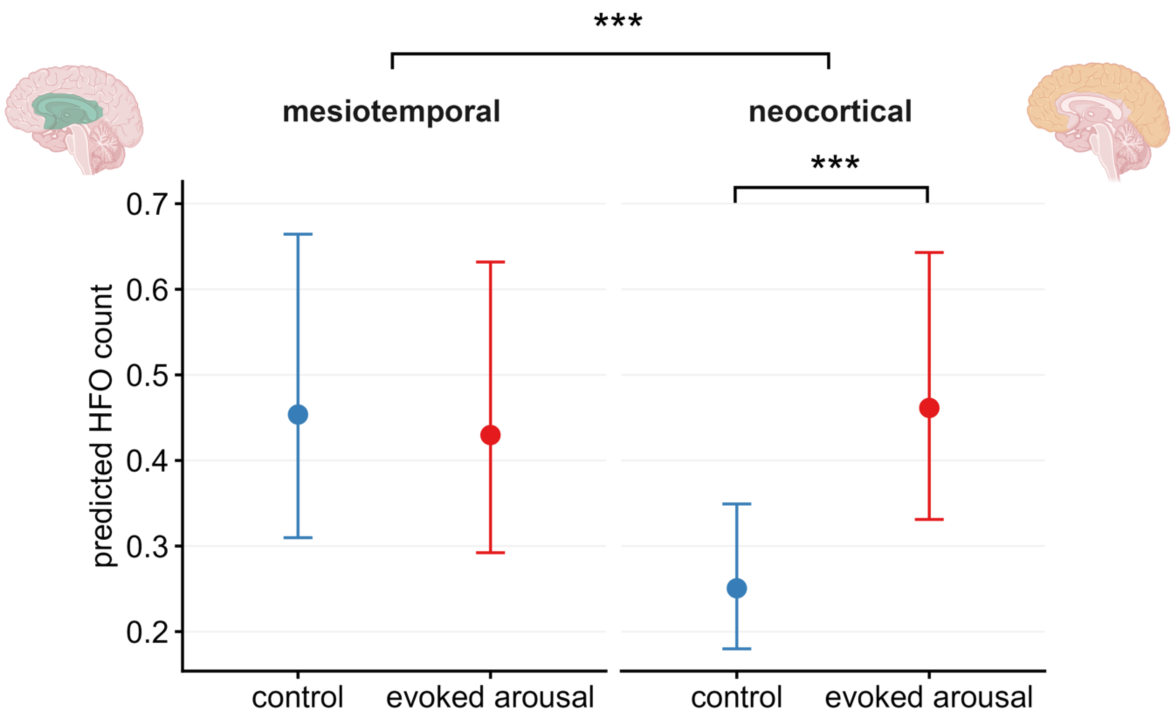


**Supplementary Figure 10. Arousal-related increases in high-frequency oscillations (HFOs) occur in neocortical but not mesiotemporal regions.** Model-predicted mean HFO counts per channel-epoch by anatomical region (mesiotemporal vs neocortical) under control and evoked arousal conditions, adjusted for pre-stimulus IED count and stimulus intensity. The interaction between condition and anatomical region was significant (generalised linear mixed-effects model (GLMM), β-estimate ± standard error (SE): 0.664 ± 0.050, incidence rate ratio (IRR) = 1.94, $p < 0.0001$), indicating that the effect of evoked arousals on HFO count differed between mesiotemporal and neocortical regions. In mesiotemporal regions ($n = 14$ patients), predicted HFO count did not differ significantly between evoked arousals and controls (GLMM, β-estimate ± SE: -0.054 ± 0.048, IRR = 0.95, $P = 0.511$). In neocortical regions ($n = 17$ patients), predicted HFO count was significantly higher during evoked arousals than controls (GLMM, β-estimate ± SE: 0.610 ± 0.015, IRR = 1.84, $p < 0.0001$), corresponding to an 84% increase. Error bars represent 95% confidence intervals. *** $p < 0.001$.

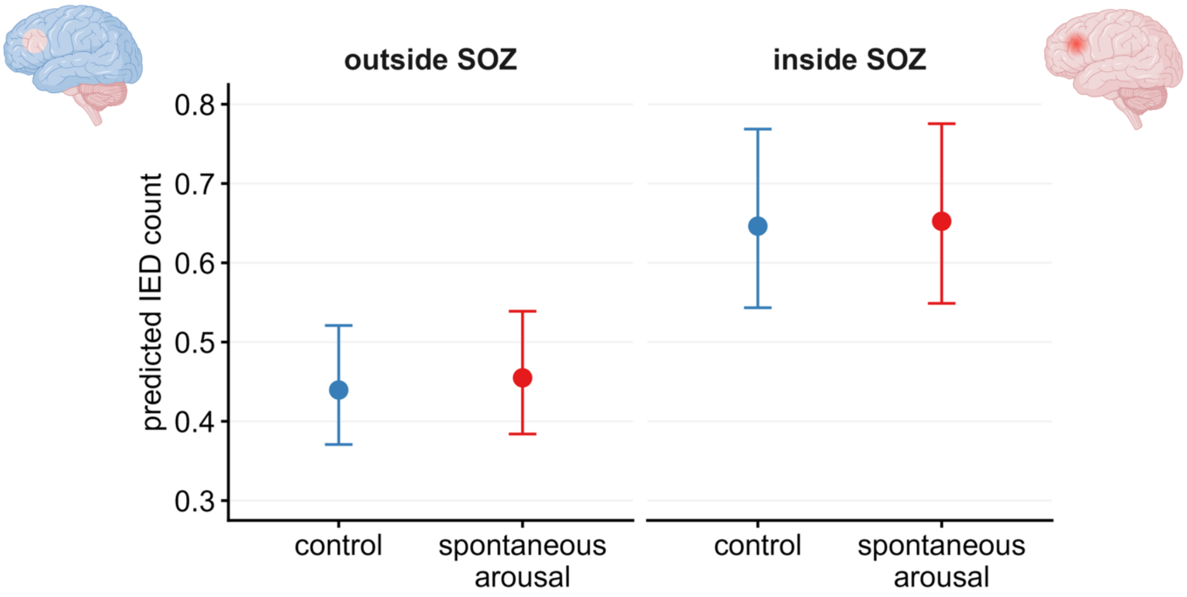


**Supplementary Figure 11. Interictal epileptiform discharge (IED) increases during spontaneous arousals are not modulated by the seizure-onset zone (SOZ).** Model-predicted mean IED counts per channel-epoch by SOZ status (outside vs inside SOZ) under control and evoked arousal conditions, adjusted for baseline IED count. The interaction between condition and SOZ status was not significant (generalised linear mixed-effects model (GLMM), β-estimate ± standard error (SE): -0.025 ± 0.021, IRR = 0.98, $P$ = 0.940), indicating that the effect of evoked arousals on IED count did not differ between contacts inside and outside the SOZ. Error bars represent 95% confidence intervals.

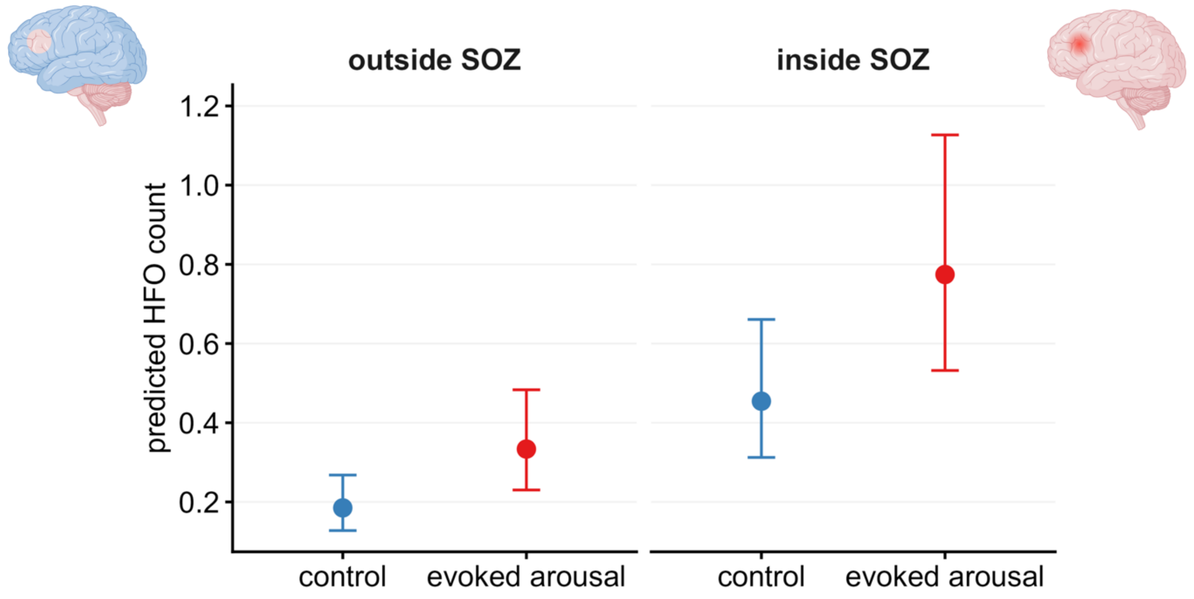


**Supplementary Figure 12. Arousal-related increases in high-frequency oscillations (HFOs) are not modulated by the seizure-onset zone (SOZ).** Model-predicted mean HFO counts per channel-epoch under control and evoked arousal conditions, adjusted for pre-stimulus HFO count and stimulus intensity. The interaction between condition and SOZ status was not significant (generalised linear mixed-effects model (GLMM), β-estimate ± standard error (SE): -0.056 ± 0.030, IRR = 0.95, $P = 0.239$), indicating that the effect of evoked arousals on HFO count did not differ between contacts inside and outside the SOZ. Error bars represent 95% confidence intervals.

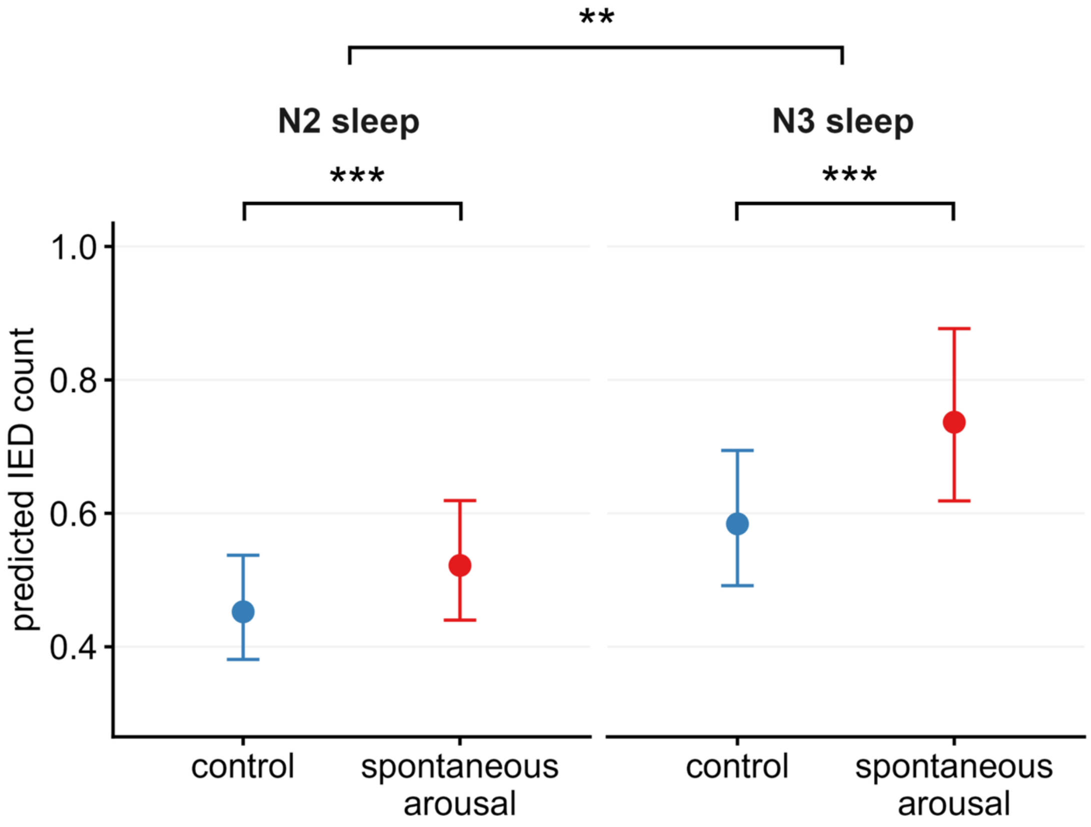


**Supplementary Figure 13. Interictal epileptiform discharge (IED) increases during spontaneous arousals are sleep-stage dependent.** Model-predicted mean IED counts per channel-epoch by sleep stage (N2 vs N3 sleep) under control and spontaneous arousal conditions, adjusted for baseline IED count. The interaction between condition and sleep stage was significant (generalised linear mixed-effects model (GLMM), β-estimate ± standard error (SE): 0.089 ± 0.029, incidence rate ratio (IRR) = 1.09, $P$ = 0.0075), indicating that the effect of spontaneous arousals on IED count differed between N2 and N3 sleep, corresponding to a 10% larger increase in N3 compared to N2. In N2 sleep ($n$ = 16 patients), predicted IED count was significantly higher during spontaneous arousals than controls (GLMM, β-estimate ± SE: 0.143 ± 0.015, IRR = 1.15, $p < 0.0001$). In N3 sleep ($n$ = 13 patients), predicted IED count was also significantly higher during spontaneous arousals than controls and with a larger magnitude than in N2 (GLMM, β-estimate ± SE: 0.232 ± 0.025, IRR = 1.26, $p < 0.0001$). Error bars represent 95% confidence intervals. ** $p < 0.01$, *** $p < 0.001$.

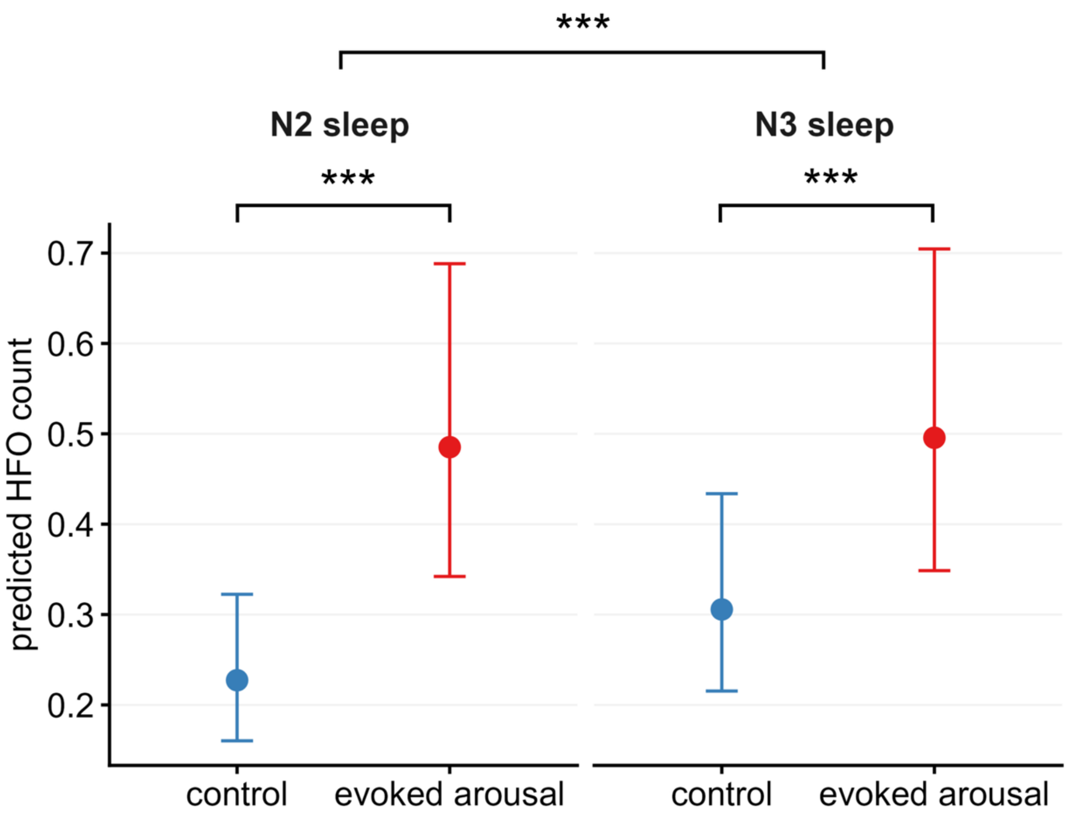


**Supplementary Figure 14. Arousal-related increases in high-frequency oscillations (HFOs) are sleep-stage dependent.** Model-predicted mean HFO counts per channel-epoch by sleep stage (N2 vs N3 sleep) under control and evoked arousal conditions, adjusted for pre-stimulus HFO count and stimulus intensity. The interaction between condition and sleep stage was significant (generalised linear mixed-effects model (GLMM), β-estimate ± standard error (SE): -0.275 ± 0.036, incidence rate ratio (IRR) = 0.76, $p < 0.0001$), indicating that the effect of evoked arousals on HFO count differed between N2 and N3 sleep, corresponding to a 24% smaller increase in N3 compared to N2. In N2 sleep ($n$ = 16 patients), predicted HFO count was significantly higher during evoked arousals than controls (GLMM, β-estimate ± SE: 0.758 ± 0.020, IRR = 2.13, $p < 0.0001$). In N3 sleep ($n$ = 13 patients), predicted HFO count was also significantly higher during evoked arousals than controls though with a smaller magnitude than in N2 (GLMM, β-estimate ± SE: 0.483 ± 0.030, IRR = 1.62, $p < 0.0001$). Error bars represent 95% confidence intervals. *** $p < 0.001$.

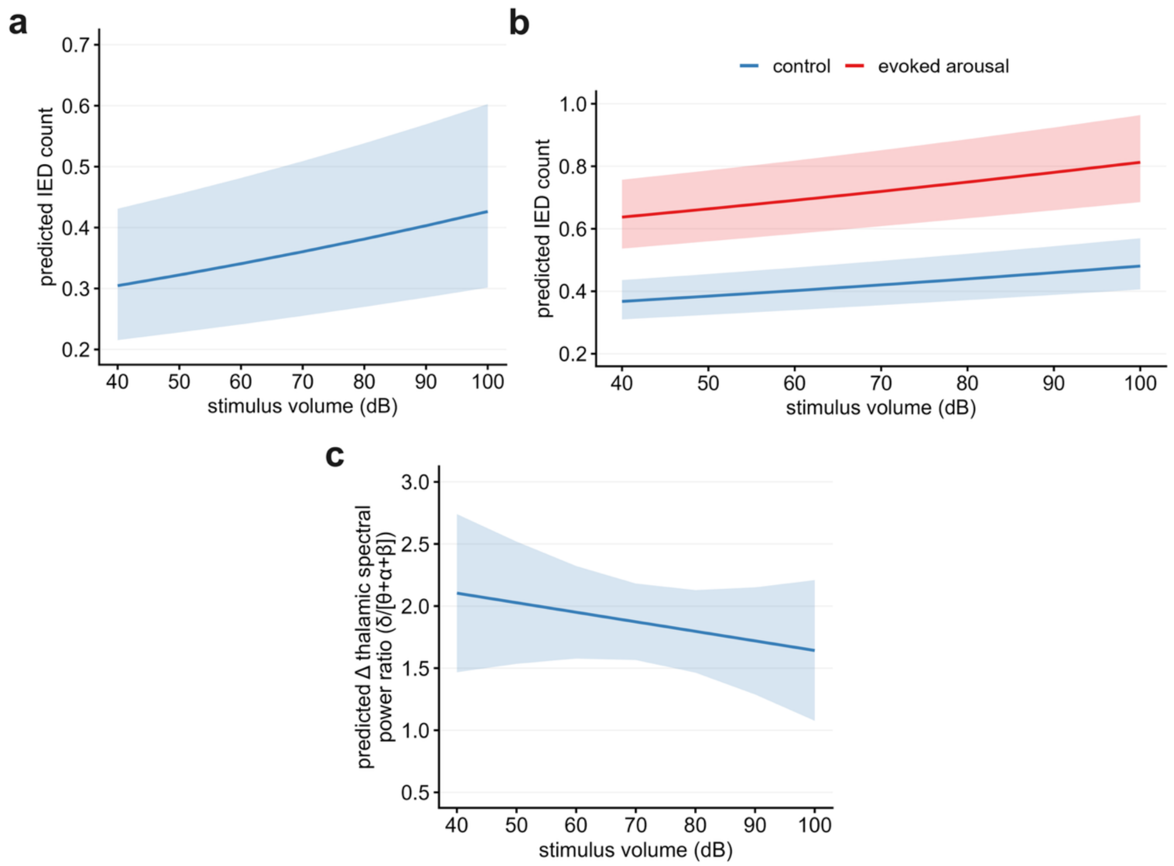


**Supplementary Figure 15. Effects of stimulus intensity on interictal epileptiform discharge (IED) counts and thalamic spectral content. (a)** Model-predicted IED count as a function of stimulus intensity (in dB) during control epochs (auditory stimuli that did not result in a detectable arousal response). Stimulus intensity was associated with a small but significant increase in IED count (generalised linear mixed-effects model (GLMM), β-estimate ± standard error (SE): 0.0056 ± 0.0005, incidence rate ratio (IRR) = 1.006, $p < 0.0001$), corresponding to a 0.6% increase in IED count per 1 dB increase in stimulus intensity. **(b)** Model-predicted IED count as a function of stimulus intensity for control epochs and evoked arousals. The interaction between event type (evoked arousal vs control) and stimulus intensity was not significant (GLMM, β-estimate ± SE: -0.0004 ± 0.0006, IRR = 1.00, $P = 0.514$), indicating that the substantially larger IED increases observed during evoked arousals did not scale with stimulus intensity and represent a distinct state change occurring independently of stimulus intensity. **(c)** Model-predicted change in thalamic spectral power ratio (δ/[θ+α+β] ([0.5–4 Hz]/[4.5–30 Hz])) as a function of stimulus intensity during control epochs, in patients with SEEG electrodes implanted within the thalamus ($n = 4$). Stimulus intensity was not significantly associated with change in thalamic spectral power ratio (GLMM, β-estimate ± SE: -0.008 ± 0.009, $P$ =

0.382), suggesting that stimulus intensity did not drive detectable shifts in thalamic spectral content in the absence of an arousal response. Shaded areas represent 95% confidence intervals.

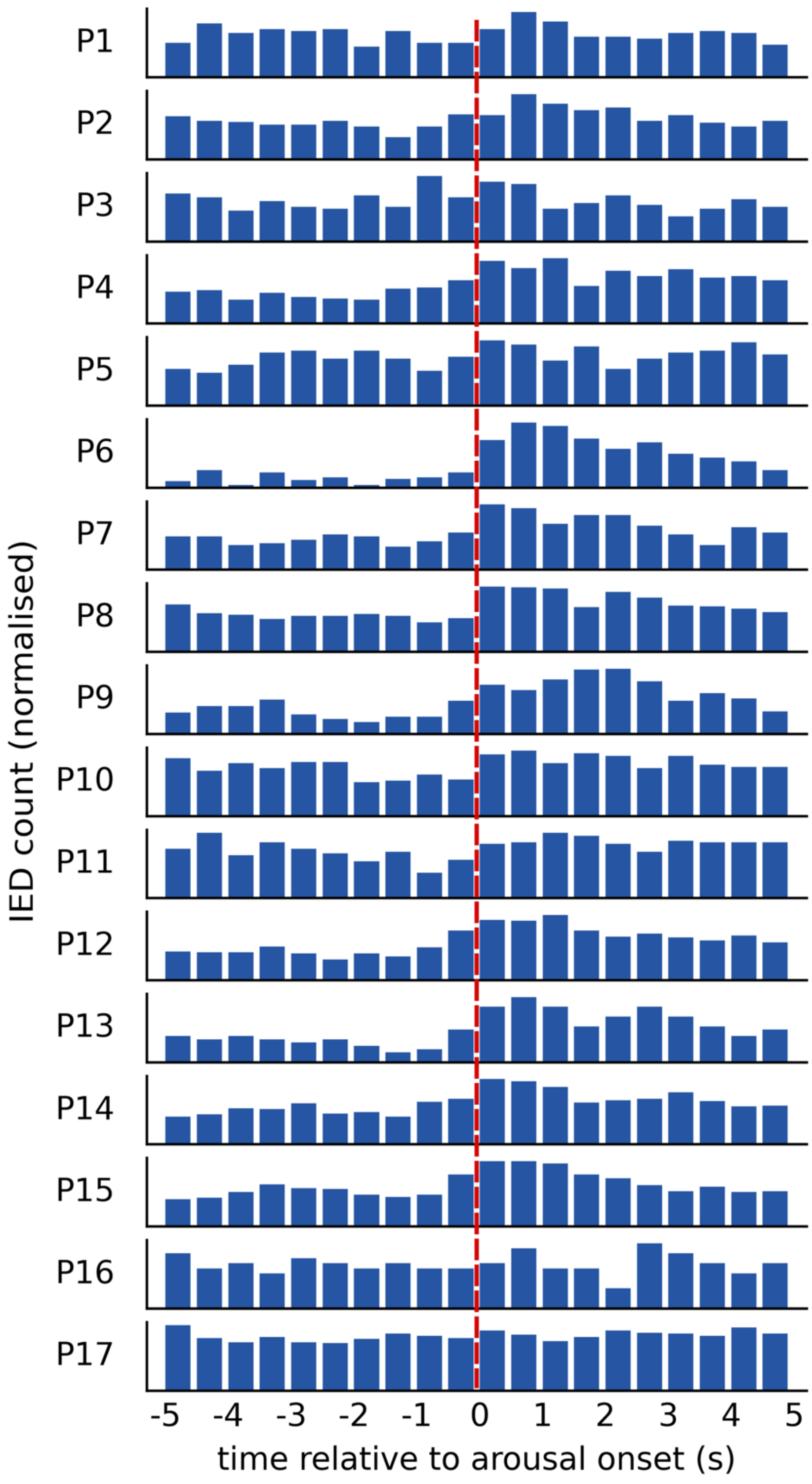


**Supplementary Figure 16. Patient-level temporal profiles of interictal epileptiform discharges (IEDs) aligned to arousal onset.** Histograms showing the temporal distribution IEDs relative to the onset of evoked arousals for each individual patient (*n* = 17), constructed using 500-ms bins. For each patient, IED counts were normalised to their maximum value, and the y-axis was uniformly scaled between 0.5 and 1 relative to this maximum to enable consistent comparison across patients. IEDs show a consistent tendency to cluster early in the arousal period, peaking shortly after the onset of evoked arousals (red dashed line) across patients, suggesting a common temporal profile of arousal-driven epileptic activity.
